# Representational drift in primary olfactory cortex

**DOI:** 10.1101/2020.09.24.312132

**Authors:** Carl E. Schoonover, Sarah N. Ohashi, Richard Axel, Andrew J.P. Fink

## Abstract

Representations of the external world in sensory cortices may define the identity of a stimulus and should therefore vary little over the life of the organism. In the olfactory system the primary olfactory cortex, piriform, is thought to determine odor identity^1–6^. We have performed electrophysiological recordings of single units maintained over weeks to examine the stability of odor representations in the mouse piriform cortex. We observed that odor representations drift over time, such that the performance of a linear classifier trained on the first recording day approaches chance levels after 32 days. Daily exposure to the same odorant slows the rate of drift, but when exposure is halted that rate increases once again. Moreover, behavioral salience does not stabilize odor representations. Continuous drift poses the question of the role of piriform in odor identification. This instability may reflect the unstructured connectivity of piriform^7–15^ and may be a property of other unstructured cortices.

## Introduction

Sensory cortices create an internal representation of the external world. If these representations provide the basis for stimulus identification, they must vary little over the life of an organism. In primary sensory neocortices tuning to basic features such as retinotopy, somatotopy and tonotopy is stable^16–26^. Responses may vary from day to day, but absent perturbation or training, this variability is bounded and differences do not accumulate over time^20,22^. In the olfactory system the specificity of odors is represented by unique and distributed ensembles of neurons in piriform cortex, and this has led to the suggestion that odor identity is established by piriform^1–6^. Sensory neurons that express the same receptor project with precision to spatially invariant glomeruli in the olfactory bulb^27^. Each odorant evokes a distinct pattern of glomerular activity that is stable over several months^28^. Axonal projections from individual glomeruli discard this spatial patterning and diffusely innervate the piriform without apparent structure^7–12^. Thus, a second transformation occurs in piriform where individual odorants activate unique, distributed and readily distinguishable ensembles of neurons^4,5,29^. Representations in piriform are therefore thought to determine the identity of an odor^1–6^. However, if piriform responses progressively change over time, piriform may not be the ultimate arbiter of odor identity posing the question of the role of primary olfactory cortex in olfactory perception.

### Longitudinal recordings in piriform

We measured the responses of neurons to a panel of neutral odorants over several weeks to assess the stability of responses in anterior piriform cortex. We developed methodology to follow the spiking of individual neurons over multiple days: to curtail damage to the tissue, a 32-site silicon probe was inserted with its shank precisely aligned to the axis of travel during implantation; to minimize movement of neurons relative to the recording sites, the probe was permanently cemented to the skull, rather than mounted on a microdrive (Fig. 1A, Extended Data Fig. 1, see Methods). We positioned the electrode sites of our silicon probe to sample superficial and deep pyramidal neurons of anterior piriform cortex layer 2/3. It is likely, however, that some of the single units isolated correspond to semilunar cells or inhibitory interneurons. This approach produces sustained signal quality, as measured by single unit yield, for at least five months, with 105 ± 29 (mean ± standard deviation) single units obtained on a given recording session (N = 97 recording sessions in 16 mice, Fig. 1B). Over this interval the number of single units isolated within a session exhibited only a slight and not significant decrease as a function of implant age (*ρ* = −0.1, p = 0.23), despite the probe remaining cemented in a fixed location with respect to the tissue (Fig. 1B). Moreover, signals were sufficiently stable as to permit isolation of single units across multiple weeks on datasets consisting of concatenated daily recordings, with a total of 379 single units held over an interval of 32 days from 6 mice (Fig. 1C).

**Figure 1.**
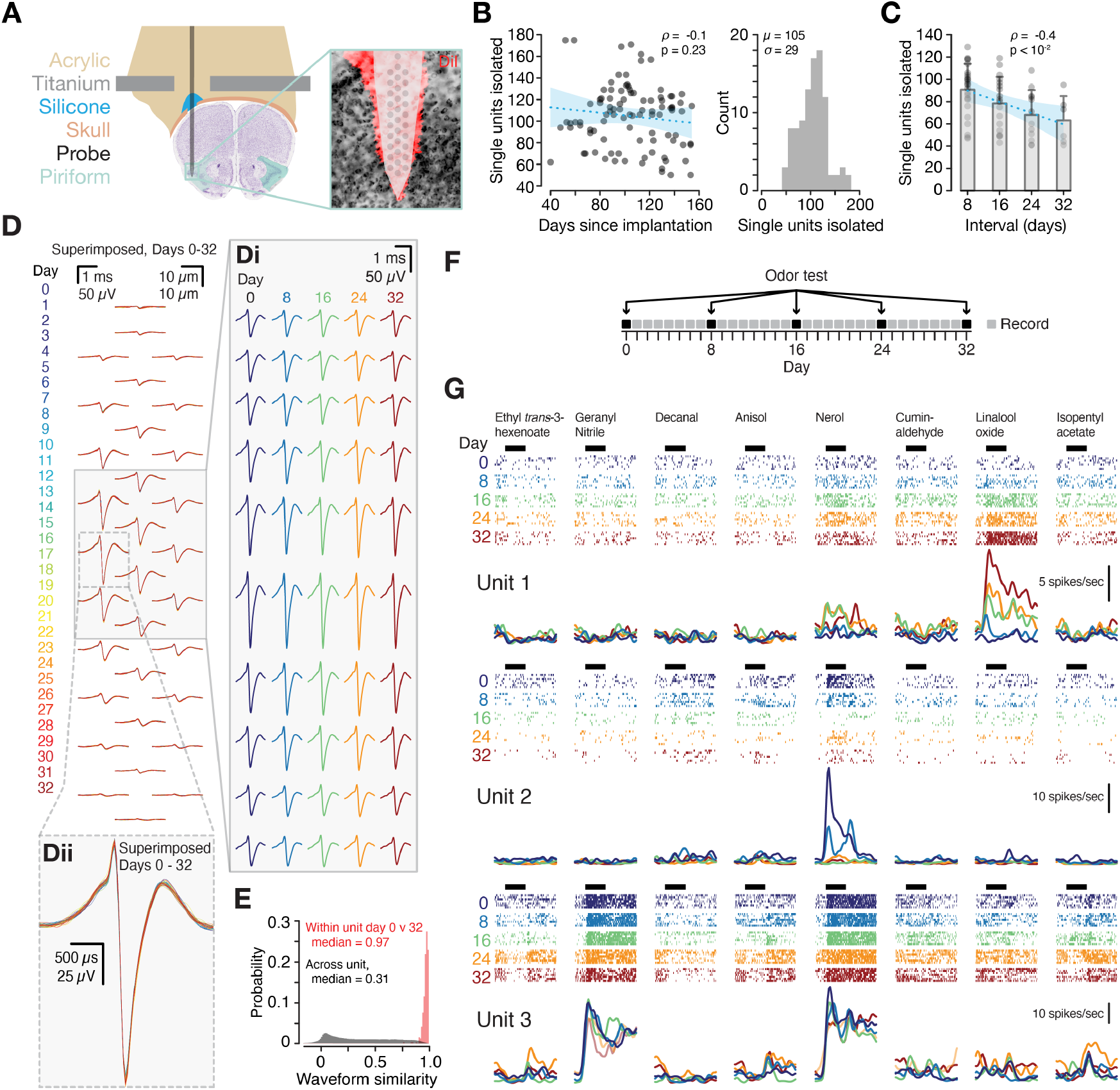
Measuring odor responses in piriform over time. **A**, Chronic silicon probe implantation in anterior piriform cortex (green). Inset: probe diagram with relative positions of the 32 recording electrodes; red, DiI marking probe position; black, cell bodies (NeuroTrace). **B**, Left, single unit yield per recording session as a function of time since probe implantation. Blue dashed line, linear regression; blue shading 95% confidence interval (C.I.), Pearson’s correlation *ρ* = −0.12, p = 0.23. Right, single unit yields across all single-day recording sessions. **C**, Number of single units retained as a function of recording interval duration. Points, individual experiments; grey bars, mean ± standard deviation; blue dashed line, linear regression; blue shading, 95% C.I., Pearson’s correlation *ρ* = −0.41, p = 1.2 × 10^−3^, N = 24 recordings across 8 days, N = 18 recordings across 16 days, N = 12 recordings across 24 days, N = 6 recordings across 32 days, all from 6 mice. **D**, Average waveforms for a representative single unit recorded at each of the 32 recording sites; each day is plotted a separate color (0-32, color scheme throughout, left), and average waveforms from all days are superimposed. Inset **Di**, mean waveforms for days 0, 8, 16, 24, and 32 for a subset of recording sites, indicated by the grey box. Inset **Dii**, mean waveforms for days 0 to 32, superimposed for a single recording site (dashed grey box). **E**, Waveform correlations for each single unit between days 0 and 32 (red) and across all single units within day 32 (black), Wilcoxon rank-sum on within-unit vs. within-day waveform correlation, p = 4.8 × 10^−246^, N = 379 single units from 6 mice. See also Extended Data Figs. 1–4 for additional single unit stability metrics. **F**, Experiment diagram. Black boxes, test odorant administration (days 0, 8, 16, 24, and 32); Grey boxes, record spontaneous activity without odorant administration. **G**, Single unit examples. Columns separate test odorants (chemical names, top). Spike rasters (rows: 7 trials per day) and peristimulus time histograms, superimposed across days, are colored by day as indicated. Horizontal black bars, 4 sec odorant pulse. See Extended Data Fig. 5 for additional single unit examples.

The shape and magnitude of spike waveforms may vary over time, confounding efforts to follow single units across days^30–32^. We monitored the consistency of both waveform and non-waveform-based features to assess the reliability of single unit tracking over time. The mean waveforms for a representative single unit recorded each day across a 32-day interval showed negligible changes in waveform shape over this period (Fig. 1D, all waveforms superimposed, Di, side-by-side, Dii, zoomin, superimposed across a 32-day interval). Moreover, waveforms were markedly stable, exhibiting high across-day correlations in waveform shape over the entire 32-day recording interval (Fig. 1E, Extended Data Fig. 1B, F, Extended Data Fig. 2B, E). The median correlation between day 0 and day 32 was 0.97 (N = 6 mice, Fig. 1E). In contrast, the correlation between a given single unit’s waveforms and those of the other single units recorded within a single day was 0.31 (Fig. 1E). Moreover, the waveforms of a given single unit were more correlated to themselves across days than to the waveforms of any other simultaneously recorded single unit on the probe (Extended Data Fig. 2B, E).

We determined whether neurons were moving relative to the probe by estimating the position of single units on each recording day. We observed a median displacement of only 3.5 μm (first quartile (Q1) = 2.1 μm, third quartile (Q3) = 5.6 μm) across the 32-day recording interval, less than half the median distance to the next most similar single unit within a given day (7.4 μm (Q1 = 4.5 μm, Q3 = 10 μm), p = 3.2 × 10^−68^, Wilcoxon rank-sum, N = 6 mice, Extended Data Fig. 2F), demonstrating negligible movement of the tissue with respect to the recording sites (Extended Data Figs. 1C, G; Extended Data Fig. 2C, F; Extended Data Fig. 3). Moreover, the shapes of single unit autocorrelograms, a feature to which the spike sorting algorithm is blind, were similarly stable across the recording period (Extended Data Fig. 1 D, H and Extended Data Fig. 2 D, G) and were not predictive of waveform similarity or displacement (Extended Data Fig. 4). Together, the stability of these features demonstrate that individual single units can be followed for extended periods.

### Drift of odor responses over time

The stability of single units over weeks permitted us to examine whether the odor responses of individual neurons are maintained over time. We recorded neural signals daily across a 32-day interval and measured responses of single units to a panel of either four or eight neutral odorants, presented seven times per day every eight days (Fig. 1F). The panel of odorant molecules was selected to maximize diversity in functional groups and organoleptic properties. Mice were awake and head fixed but were not engaged in any task other than sampling the odor stimuli that were presented.

We observed gradual and pronounced changes in odor responses over time. Single units either gained (Fig. 1G, top) or lost (Fig. 1G, middle) responsivity to a given odorant and only on rare occasions exhibited stable responses to the odorant panel (2.8% of single units) over 32 days (Fig.1G, bottom). The progressive changes in the responses of single units were not due to a global loss of responsiveness in piriform, but rather to continual alterations in odor representations (Fig. 2A). We quantified this drift by comparing the response magnitude of odor-unit pairs across days and found that responses became increasingly dissimilar over time (Fig. 2B, within-day R^2^ = 0.94, 8-day interval R^2^ = 0.52, 16-day interval R^2^ = 0.31, 24-day interval R^2^ = 0.22, 32-day interval R^2^ = 0.08, N = 6 mice). In contrast, changes in spontaneous firing rates across the same intervals were relatively small (Extended Data Fig. 6). Moreover, the performance of a linear classifier trained on earlier days and tested on later days deteriorated as a function of time between training and testing, approaching chance levels after 32 days (Fig. 2C). This drift was symmetric in time (Extended Data Fig. 7A) and classification accuracy was not improved by concatenating multiple days in the training set (Extended Data Fig. 7B). Thus, it is not possible to establish single units most informative about odor identity on day 32 based on their responses across days 0 through 24. These data indicate that changes in odor representations in piriform accumulate over time.

**Figure 2.**
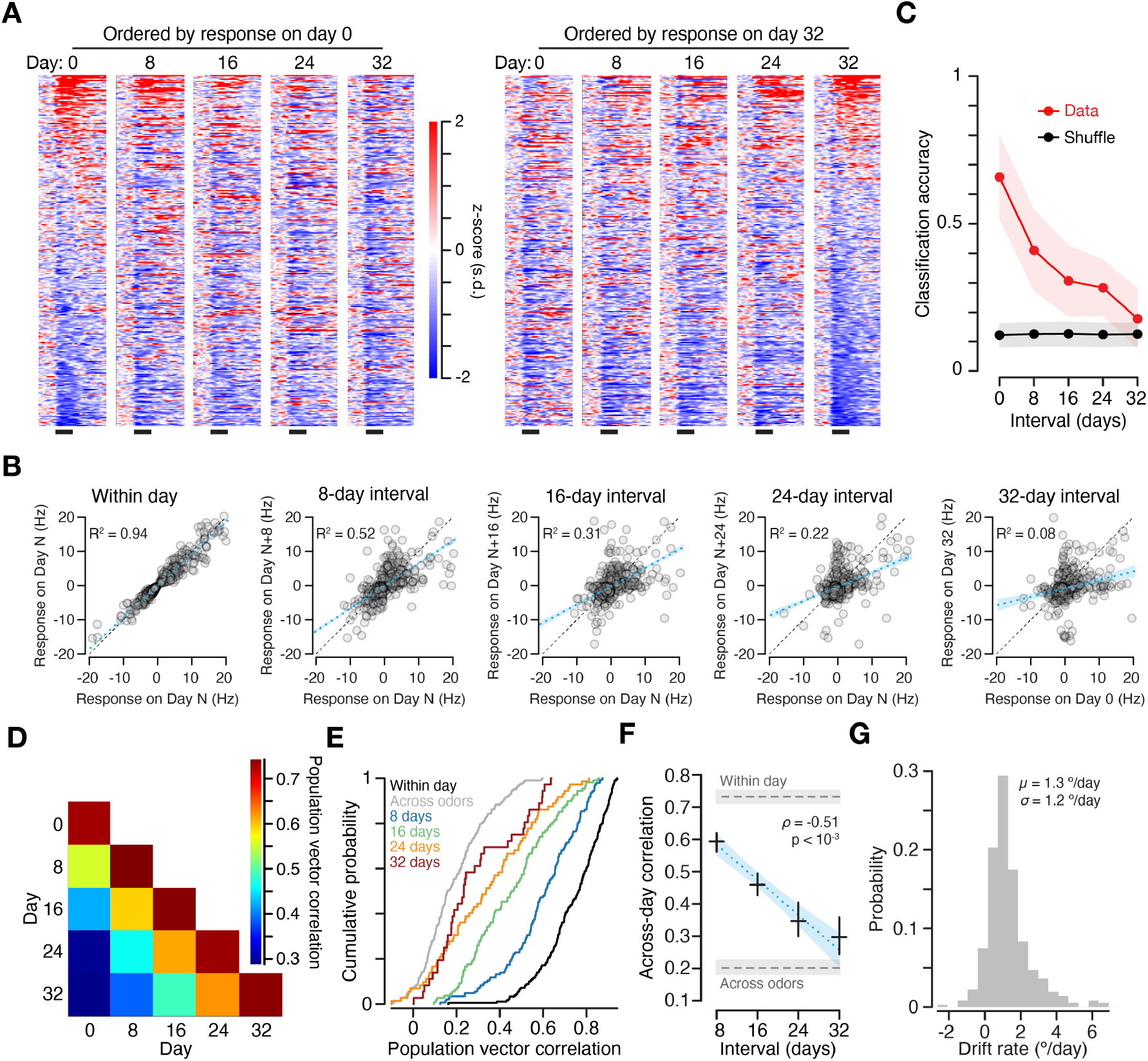
Drifting odor representations in piriform. **A**, Odor response heat maps for 300 randomly selected odor-unit pairs, ordered by response magnitude on day 0 (left five columns) and response magnitude on day 32 (right five columns). Responses are *z*-scored and plotted in units of standard deviation (s.d.). Black bars, 4 sec odorant pulse. **B**, Mean spontaneous baseline-subtracted evoked responses of odor-unit pairs within a single day (far left, N = 2,353 odor-unit pairs) and across intervals of 8 days (N = 2,843 odor-unit pairs), 16 days (N = 1,879 odor-unit pairs), 24 days (N = 1,170 odor-unit pairs), and 32 days (N = 577 odor-unit pairs). Data pooled across 6 mice. Within-day, trial-averaged responses on odd trials plotted against trial-averaged responses on even trials. Across-day, response magnitude on later days plotted against response magnitude on earlier days. Black dashed line, unity; blue dashed line, linear regression; blue shading, 95% C.I. Regression for each interval was performed across all odor-unit pairs that showed a significantly modulated response on at least one of the two days that are compared (Wilcoxon rank-sum, *α* = 0.001). Each plot shows a random subset of 577 odor-unit pairs, to match the number of significant odor-unit pairs measured across the 32-day interval (right panel). **C**, Red, classification accuracy (support vector machine, linear kernel) of single-trial population vectors as a function of interval (randomly selected subsets of 41 single units each, 8-way classification, from the N = 3 animals that were presented a test odorant panel of 8 stimuli). Classification performance on day 0 computed using leave-one-out cross-validation. For all other intervals the model is trained on all responses from the earlier day and tested on all responses from the later day. Black performance with stimulus labels shuffled. Points, mean; Shading, standard deviation. **D**, Pearson’s correlations of trial-averaged population vectors, averaged across all test odors. Within-day correlations measured between trial-averaged population vectors on even versus odd trials. **E**, Cumulative distribution function of trial-averaged population vector correlations. Black, cumulative distribution of within-day correlations; Grey, cumulative distribution of across-odor correlations. **F**, Mean across-day correlations (black crosses) and 95% bootstrap confidence interval of the mean (95% C.I.) for pairs of trial-averaged population vectors in 6 mice: 8-day interval, N = 144 pairs; 16-day interval, N = 108 pairs; 24-day interval, N = 72 pairs; 32-day interval, N = 36 pairs; within day, all intervals N = 360 pairs. Blue dashed line, linear regression; Blue shading, 95% C.I., *ρ* = −0.51, p = 1.2 × 10^−24^; Grey dashed line, top, mean within-day correlations and 95% C.I.; Grey dashed line, bottom, mean across-odor correlations and 95% C.I. **G**, Distribution of drift rates measured across all test odors and intervals in 6 mice, N = 360 pairs of trial-averaged population vectors. See Extended Data Fig. 8 for calculation of drift rate.

We next estimated the rate of drift of odor representations in piriform. First we computed the correlation between trial-averaged population odor responses across all pairs of recording days (Fig. 2D-F). Consistent with across-day changes in single unit response magnitude (Fig. 2B), population vector correlations across days decreased as a function of interval (Fig. 2F). We then used these correlations to compute the angle between trial-averaged population vectors (Extended Data Fig. 8). An exponential fit to these measurements gave a time constant of 28.3 days with an asymptote of 89.4° (an angle of 90° indicates complete decorrelation, Extended Data Fig. 8B, right). We estimated the rate of drift in two ways. First, we computed the rate of change of the exponential fit over the 32-day interval, finding a mean rate of 1.0 °/day. Second, we estimated the drift rate between every pair of days. The angle between days reflects a combination of both across-day drift in odor representations and within-day trial-to-trial variability^33^. We therefore subtracted within-day variability from across-day variability and divided by time (Extended Data Fig. 8A). This estimate produced a drift rate of 1.3 ± 1.2 °/day across all intervals (mean ± s. d., N = 6 mice, Fig. 2G), in accordance with the estimate based on the exponential fit. Thus, piriform exhibits representational drift: changes in responses accrete continuously over time and trend towards complete decorrelation.

The changes in odor representations we have observed could be attributed to gross changes in the population response. We found, however, that the basic response properties of the piriform population change only marginally over time (Fig. 3). The fraction of single units responsive to an odor stimulus was largely unaltered across the 32-day interval (Fig. 3B, R^2^ = 8.0 × 10^−5^). Moreover, lifetime sparseness (R^2^ = 4.4 × 10^−3^), population sparseness (R^2^ = 3.1 × 10^−2^), within-day classifier performance (R^2^ = 5.2 × 10^−2^), and within-day variability (R^2^ = 5.3 × 10^−5^) varied little (Fig. 3B). We note that within-day classifier performance exhibited modest deterioration (Fig. 3B, second from right), which may contribute a small fraction of the decrease in across-day classification accuracy we measure over time (Fig. 2C). These observations demonstrate that drift in odor representations is not the result of a global diminution in responsivity and occurs against a background of grossly stable population response properties.

**Figure 3.**
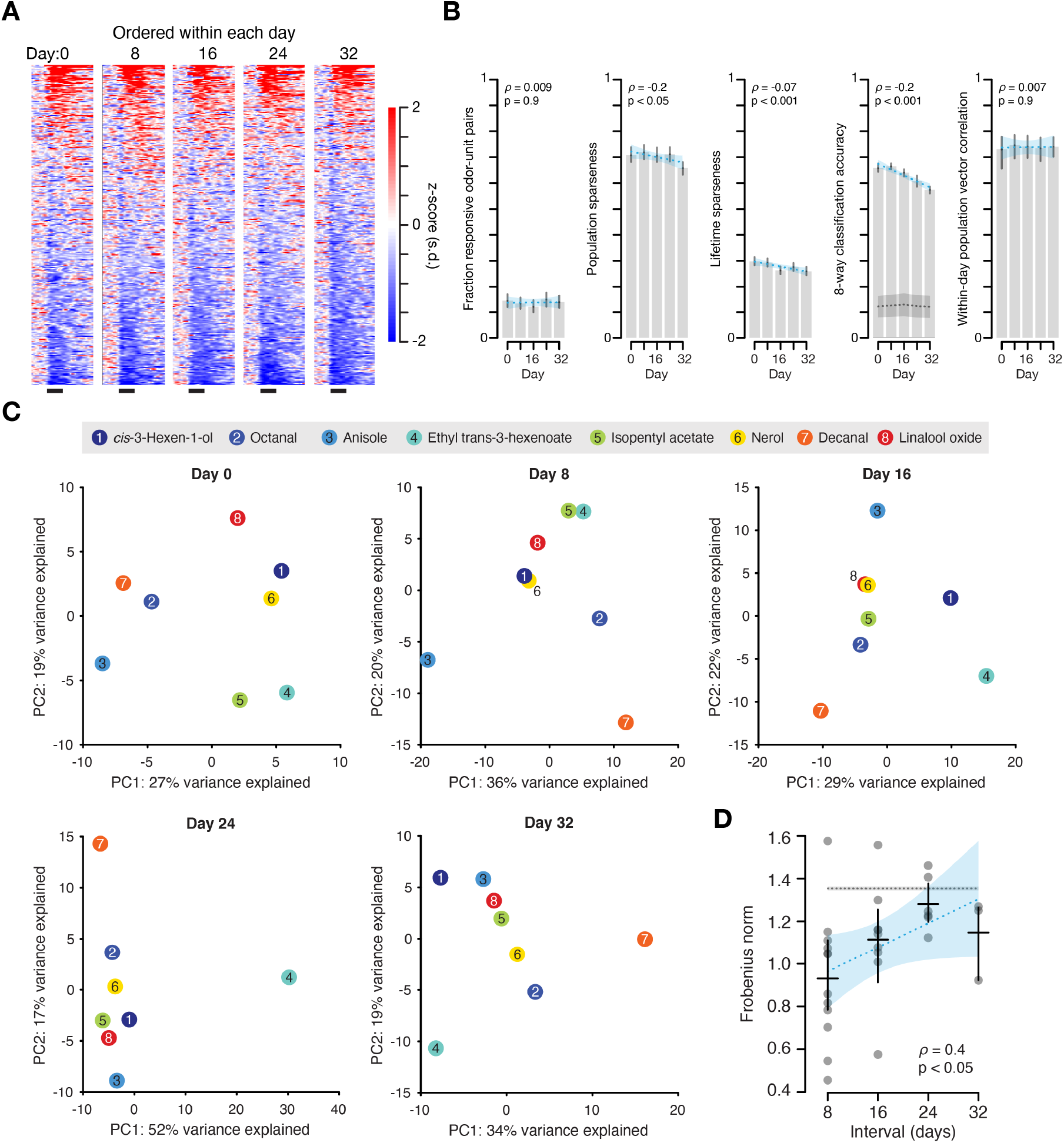
Drifting population response geometry despite stable response statistics. **A**, Odor response heat maps for the randomly selected odor-unit pairs shown in Fig. 2A, but ordered by response magnitude on each day. Responses are *z*-scored and plotted in units of standard deviation (s.d.). Black bars, 4 sec odorant pulse. **B**, Population statistics across days. Left: *fraction of odor-unit pairs showing significant responses*, Wilcoxon rank-sum test, *α* = 0.001, 36 animal-odor pairs for each day. Second from left: *population sparseness*. Middle: *lifetime sparseness*. Second from right: *within-day leave-one-out cross-validated classification accuracy*, support vector machine, linear kernel; in 3 animals that were presented a test odorant panel of 8 stimuli, 41 single units in each mouse; black dashed line mean and shading standard deviation, shuffled stimulus labels. Right: *within-day trial-averaged population vector correlation* for even vs. odd trials: 72 trial-averaged population vector pairs for each day from 6 mice. Throughout: grey bars, mean; black lines, 95% C.I.; red dashed line, linear regression; blue shading, 95% C.I. Estimates of fraction of responsive odor-unit pairs, population and lifetime sparseness were made using: day 0, N = 584 single units; day 8, N = 635 single units; day 16, N = 629 single units; day 24, N = 593 single units; day 32 N = 545 single units from 6 mice. **C**, Projection onto the first two principal components of trial-averaged population vectors measured in response to eight odorant stimuli in one animal. The principal components were computed separately on each one of the five days. Stimulus identity is denoted by circle color and number. **D**, Frobenius norm between pairs of odor-odor correlation matrices. 8×8 correlation matrices were computed based on trial-averaged population vectors to eight odorant stimuli recorded on each of the five days in the three animals presented a panel of 8 odorants. Frobenius norm day 8, N = 12 pairs; day 16, N = 9 pairs; day 24, N = 6 pairs; day 32, N = 3 pairs; shuffle, N = 3,000 pairs. Blue dashed line, linear regression (*ρ* = 0.4, p = 0.023); Blue shading, 95% C.I.; Grey dashed line, top, mean Frobenius norm computed using shuffled stimulus identities, grey shading, 95% C.I.

Regions downstream of piriform could in principle compensate for drift if the geometry of population representations were conserved over time, in spite of day-to-day changes in the set of neurons that respond to a given odorant. It has recently been shown that the correlation structure of piriform representations of highly similar odorant molecules is conserved across individuals, but that no such structure is apparent for dissimilar stimuli such as the ones we employed^34^. Accordingly, when we visualized a low dimensional projection of population responses onto the first two principal components (computed separately for each day and explaining between 46% and 69% of the variance), similarities between odor responses did not appear to be preserved over time (Fig. 3C). We quantified the extent to which odor response geometry is conserved by first computing the correlation matrix of population responses for each day and then computing the Frobenius norm between pairs of correlation matrices, where 0 indicates perfectly preserved relationships between odor representations and increasing values indicate increasingly dissimilar geometries. We found that the Frobenius norm rose significantly as a function of interval between the measurements and approached the mean Frobenius norm between correlation matrices computed using shuffled stimulus labels (*ρ* = 0.4, p = 0.023, Fig. 3D). Thus, the geometry of population representations of highly dissimilar odorant molecules is not conserved, but rather changes gradually. This analysis does not require the longitudinal observation of individual neurons across days, providing indirect confirmation that odor representations in piriform change over time.

It has been suggested that activity in piriform during the early phase of the odor response epoch is sufficient to accurately establish its identity^5,6,35,36^. We therefore asked if the odor response in piriform during the 200 milliseconds after the first sniff provides a stable representation of odorant stimuli. We trained a linear classifier using spikes recorded during this early epoch and then tested the performance of that classifier on data obtained on subsequent days during the same epoch. We found that classification accuracy during the initial phase of the odor response was not stable over time but rather deteriorated across days from 0.74 ± 0.12 on day 0 to 0.33 ± 0.10 on day 32 (mean ± s. d., chance = 0.25; Extended Data Fig. 9).

Finally, variance in single unit waveform stability metrics did not explain variance in changes of odor responses (Extended Data Fig. 1B-D, bottom and Extended Data Fig. 1F-H, bottom). This suggests that unstable tracking of single units across days cannot account for changes in the odor responses of single units over time. Moreover, given that these changes in odor responses did not vary around a constant mean value but rather accumulated across days, we can rule out inconsistency in stimulus delivery across test days, as well as changes in the animals’ internal state (e.g. arousal) as the cause of the representational drift.

### Frequent experience, but not salience, reduces the rate of drift

We next asked whether the representation of a behaviorally salient odorant might exhibit greater stability. We placed implanted mice in a conditioning chamber and presented conditioned stimuli: one paired with foot shock (CS+) and one presented without shock (CS−) eight times each over the course of a single session (Fig. 4A). The following day we used a Virtual Burrow Assay^37^ to confirm that animals responded selectively to the CS+, but not to the CS− or to other odorants that were never presented in the conditioning chamber. Animals showed selective ingress (escape) to the CS+ across the entire duration of the protocol (Fig. 4B). Yet, simultaneously acquired recordings showed that the representations of conditioned stimuli drifted at a comparable rate to that of neutral odors (Fig. 4C, mean (95% C.I.), CS+: 1.2 (0.7 - 1.8) °/day, CS−: 0.9 (0.7 - 1.7) °/day, Neutral: 1.0 (0.8 - 1.0) °/day, for all comparisons p ≥ 0.61, Wilcoxon rank-sum, N = 5 mice). Thus, behavioral salience does not appear to stabilize odor representations in piriform: we observe stable behavior in spite of drifting representations.

**Figure 4.**
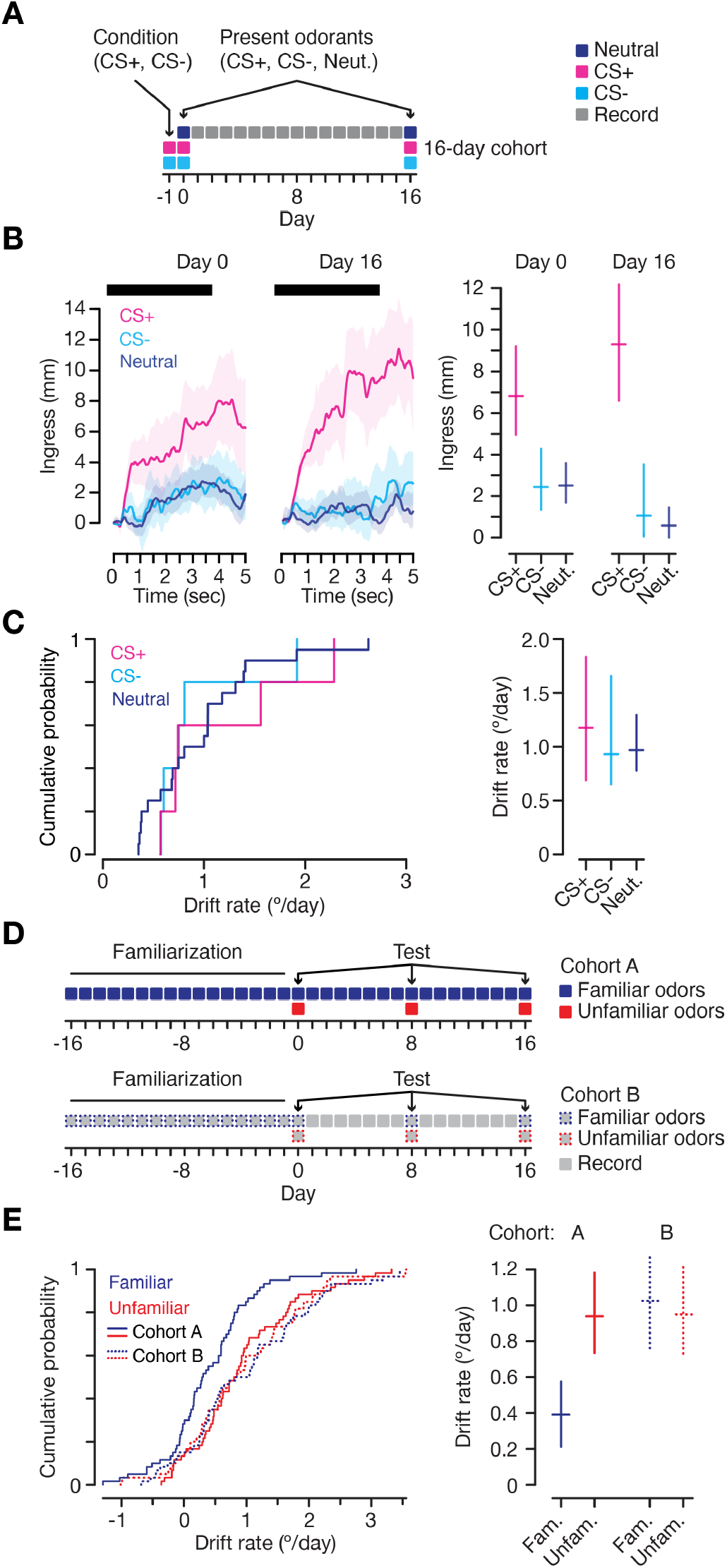
Frequent experience, but not salience, reduces the rate of drift. **A**, Conditioning experiment diagram. Magenta and cyan squares, day -1: present one odorant paired with shock (CS+) and a second without shock (CS−) in a conditioning chamber. Magenta, cyan and dark blue squares, days 0 and 16: administer conditioned and neutral odorants to head fixed animal while recording neural signals and measuring behavioral responses in Virtual Burrow Assay^37^. Grey squares, record neural signals in head fixed animal. **B**, Behavior. Left, trial-averaged (N = 5 mice) ingress amplitude across time on days 0 and 16 on trial blocks 2-7 (shading 95% C.I.). Black bar, 4 sec odorant stimulus. Right, mean (95% C.I.) ingress amplitude during the final second of the odorant epoch on blocks 2-7. For days 0 and 16, CS+ vs. CS− and CS+ vs. Neutral, p < 1.4 × 10^−3^, Wilcoxon rank-sum. **C**, Neurophysiology. Left, cumulative distributions and right, mean ± 95% C.I. (N = 5 mice) of drift rates for conditioned (CS+, magenta and CS−, cyan) and neutral (dark blue) odors. **D**, Experiment diagram. Solid blue squares: Two cohorts of mice were presented a panel of four neutral odorants daily across a 16-day interval (days -16 to 0), after which cohort A (N = 5 mice) continued to be presented the familiar odorants daily across a second 16-day interval (days 0 to 16, solid blue squares), whereas cohort B (N = 5 mice) encountered the familiar odorants at only 8-day intervals across the same period (dashed blue squares). Starting on day 0 a panel of unfamiliar odorants (solid and dashed red squares) was presented at 8-day intervals to both cohorts. In both cohorts neural signals were recorded daily regardless of whether odorants were presented. **E**, Left, cumulative distributions of drift rates measured for familiar (solid blue curve) and unfamiliar odors (solid red curve) for cohort A and for familiar (dashed blue) and unfamiliar (dashed red) odors for cohort B. Right, mean drift rate and 95% C.I. for both familiar and unfamiliar odors in cohorts A and B (drift rate familiar odors for cohort A vs. all others, p < 9.4 × 10^−4^, Wilcoxon rank-sum; N = 60 trial-averaged population vector pairs.)

Representational drift is a hallmark of a learning system that continuously updates and overwrites itself^38^. Two observations support the view that piriform operates as such a learning system. We presented a panel of odorants daily across a 32-day interval. At day 16 we also presented a set of unfamiliar odorants at 8-day intervals and compared the stability of the representations of the familiar and unfamiliar odors (Fig. 4D, top). We found that the rate of drift of the representations of familiar odors was less than half that of unfamiliar odors (Fig. 4E, solid lines, familiar odors: 0.4 (0.2 - 0.6) °/day vs. unfamiliar odors: 0.9 (0.7 - 1.2) °/day, p = 4.3 × 10^−4^, Wilcoxon rank-sum, N = 5 mice). Thus, the drift rate of piriform representations exhibits history dependence: continual experience with an odorant enhances the stability of its representation.

Importantly, this observation provides independent evidence for our ability to follow a fixed population of single units over time. We observed rapidly drifting representations for unfamiliar odors and slowly drifting representations for familiar odors, on separate trials within the same recording session of the same population of neurons. Thus, it is unlikely that the changes in odor representations we have observed are caused by a failure to follow the same population of neurons.

Models predict that in a highly plastic learning system the stable representations that encode memories can be overwritten unless the circuit has a mechanism to store them for the long term^38^. We therefore asked whether piriform has a mechanism to retain a stabilized representation after daily experience ceases. We presented a panel of odorants every day for sixteen days. This panel, along with a set of unfamiliar odorants, were then presented at 8-day intervals (Fig. 4D, bottom). We observed that once daily odor exposure ceases, representations of the familiar and unfamiliar stimuli drifted at comparable rates (Fig. 4E, dashed lines, familiar odors: 1.0 (0.8 - 1.3) °/day vs. unfamiliar odors: 1.0 (0.7 - 1.2) °/day, p = 0.81, Wilcoxon rank-sum, N = 5 mice). These data demonstrate that daily exposure slows representational drift, but without ongoing exposure drift increases once again.

## Discussion

Stimulus representations in piriform exhibit rapid and cumulative reorganization over time. Perceptual constancy, however, requires a stable source of information about the nature of the sensory stimulus. Therefore, primary olfactory cortex may not be the ultimate arbiter of the identity of odors over the life of an organism. This poses the question of the role of piriform in olfactory perception.

Piriform exhibits representational drift, history-dependent stabilization, and subsequent drift of previously stabilized representations. These data are consistent with a model in which piriform functions as a fast learning system that continually learns and continually overwrites itself. Piriform may operate as a ‘scratch pad’, rapidly encoding memory traces upon limited odorant exposure; however, it lacks a mechanism to stabilize these memory traces and therefore cannot store those memories over the long term^38^. This may explain why the representation of an odor changes when an animal isn’t experiencing it. In a fast learning system, experience with one odorant will drive plastic changes that alter the representations of other odors. In our experiments ongoing experience with the odorants of the homecage, for example, may result in cumulative changes in the representations of our test odors. In this model representational drift in piriform is a consequence of continual learning and concomitant overwriting.

Learned representations in piriform, reflective of recent statistics of the olfactory environment, may be only transiently useful to the animal since they are continuously updated as the environment changes. Alternatively, representations in piriform may be transferred to a more stable region downstream for long-term storage. Models that posit the consolidation of memory traces from a highly plastic, unstable network to a less plastic, stable network computationally outperform single-stage learning systems by combining both flexibility and long-term memory^39^. This framework has been proposed to account for the initial formation of episodic memories in hippocampus and later consolidation in neocortex^40,41^. Piriform may therefore operate in conjunction with a slow learning system downstream to support long-term access to stored representations. This hypothetical slow learning system must receive input from olfactory bulb where representations are stable^28^, as well as from piriform, where they are drifting. This third region may, in concert with piriform, accommodate the identification of an olfactory stimulus.

Alternatively, regions downstream of piriform could in principle compensate for drift independent of a stable downstream structure, either by continuous updating maintained by offline replay^42^, or by leveraging some invariant geometry in the population representation in piriform^43,44^. The latter is unlikely considering our observation that the representational geometry itself drifts over time. Finally, these observations do not rule out that piriform represents odor identity along with other unknown variables in a format not readily captured by simply examining responses to odor stimuli over time^45^.

It is possible, however, that primary olfactory cortex does not play a role in establishing the identity of an odor. Piriform is not the sole recipient of olfactory sensory information. Stable representations of odor in the olfactory bulb are broadcast to diverse regions that link sensation with valence and action: the anterior olfactory nucleus, olfactory tubercle, cortical amygdala and lateral entorhinal cortex^7,46^

The stability of neural representations varies across brain regions. Representational drift has been previously reported in hippocampus CA1^47–50^ and posterior parietal cortex^51^. In primary sensory neocortices, however, absent a training paradigm or a gross perturbation, responses vary across days but tuning remains centered around a fixed mean^20,22,23,26,30,52–54^. Moreover, tuning changes induced in sensory neocortex by training or perturbation have been found to reverse after the intervention is discontinued^20,24^. In contrast, our data show that in primary olfactory cortex changes in odor representations increase continuously: the population representation at every successive time point is increasingly dissimilar to the first.

What conditions promote drifting versus stable representations? The anatomical organization of most sensory cortices may set a bound on changes in stimulus tuning to basic features. In primary visual cortex, for example, inputs from thalamus obey ordered retinotopic organization^55^. The weakening or loss of one synapse onto a visual cortical neuron is therefore likely to be matched by the strengthening or gain of a synapse tuned to a similar retinotopic region, limiting representational drift. However, the organization of piriform differs from sensory neocortex: piriform receives unstructured inputs from olfactory bulb^7–12^, and recurrent cortical connections are broadly distributed and lack topographic organization^13–15^. Representational drift in piriform may be explained by ongoing plasticity in an unstructured network. The structured connectivity observed in most early sensory regions may not be present in higher centers, suggesting that representational drift may be a pervasive property of cortex.

## Methods Summary

Procedures performed in this study were conducted according to US National Institutes of Health guidelines for animal research and were approved by the Institutional Animal Care and Use Committee of Columbia University. See **Full Methods**.

## Acknowledgements

We are grateful to L.F. Abbott, S.R. Datta, S. Fusi, J.W. Krakauer, A. Kumar, M.A. Long, A.M. Michaiel and S.L Pashkovski for comments on the manuscript; G.W. Johnson and T. Tabachnik for assistance with instrumentation; G. Buzsáki and members of the Buzsáki laboratory, D.A. Gutnisky, and C.C. Rodgers for experimental advice; the instructors of the 2015 Advanced Course in Computational Neuroscience at the Champalimaud Foundation for guidance; D.F. Albeanu and members of the Albeanu laboratory, D. Aronov, K.A. Bolding, R. Costa, J.P. Cunningham, J.T. Dudman, W.M. Fischler, K.M. Franks, M.E. Hasselmo, V. Jayaraman, A.Y. Karpova, E. Marder, J.A. Miri, C. Poo, D. Rinberg and members of the Rinberg laboratory, and E.S. Schaffer for insightful comments; M. Gutierrez, C.H. Eccard, P.J. Kisloff, and A. Nemes for general laboratory support; and the Howard Hughes Medical Institute and the Helen Hay Whitney Foundation for financial support.

## Author contributions

This work is the result of a close collaboration between C.E.S. and A.J.P.F., who conceived the study and designed the experiments. Experiments were performed by C.E.S., S.N.O., and A.J.P.F. The data were analyzed and the manuscript was written by C.E.S, R.A., and A.J.P.F.

## Data availability

Data will be made available upon reasonable request.

## Code availability

Code will be made available upon reasonable request.

---

**Extended Data Figure 1.**
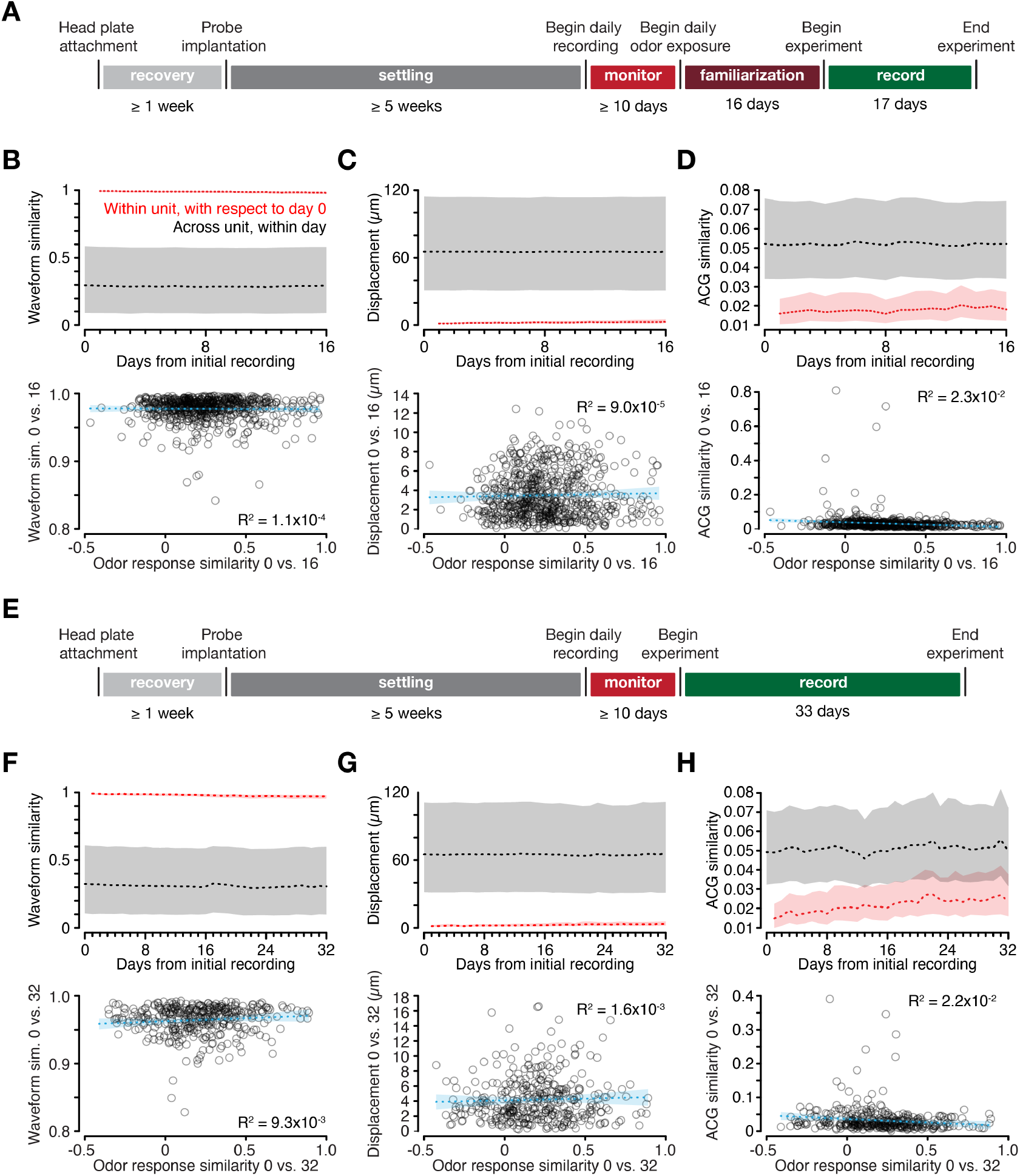
Quantifying single unit stability across days. **A**, Experiment time course for 17-day experiments. *Recovery*: minimum two week period after headplate attachment and stereotactic targeting prior to silicon probe implantation to allow full recovery; *Settling*: minimum five week period after probe implantation to permit tissue settling and signal stabilization; *Monitor*: minimum 10-day period during which neural signals were recorded daily to assess signal stability. Experiments only commenced once single units could be reliably tracked across days; *Familiarization*: daily odorant presentation for experiments described in Fig. 4D-E; *Record*: 17-day recording period during which odorant was administered and drift rate was measured. **B**, Top, for single units held across days, Pearson’s correlation between waveforms (waveform similarity) measured on day 0 and waveforms measured on subsequent days (red) and across all single units within each day (black). Dashed line, median. Shading, indicates boundaries of top and bottom quartiles (for all 17-day experiments, N = 690 single units from 7 mice). Bottom, Black circles, Pearson’s correlation of single unit mean spike waveforms measured between days 0 and 16 and plotted against the Pearson’s correlation of the vector of trial-averaged responses of a given single unit to each odorant of a panel; Blue dashed line, linear regression; Blue shading, 95% C.I. **C** and **D**, Same analysis as **B** but for single unit centroid displacement and similarity (Euclidean norm) of the shape of single unit spike time autocorrelograms (ACG), respectively. **C**, bottom, single unit centroid displacement vs. odor response similarity and **D**, bottom, spike time ACG similarity vs. odor response similarity. **E-H** is the same as **A-D** but for 33-day experiments (for all 33-day experiments, N = 379 single units from 6 mice). **F**, top, waveform similarity across 33 days. Bottom, waveform similarity vs. odor response similarity. **G**, top, centroid displacement across 33 days. Bottom, single unit centroid displacement vs. odor response similarity and **H**, top, ACG similarity across days. Bottom, spike time ACG similarity vs. odor response similarity.

**Extended Data Figure 2.**
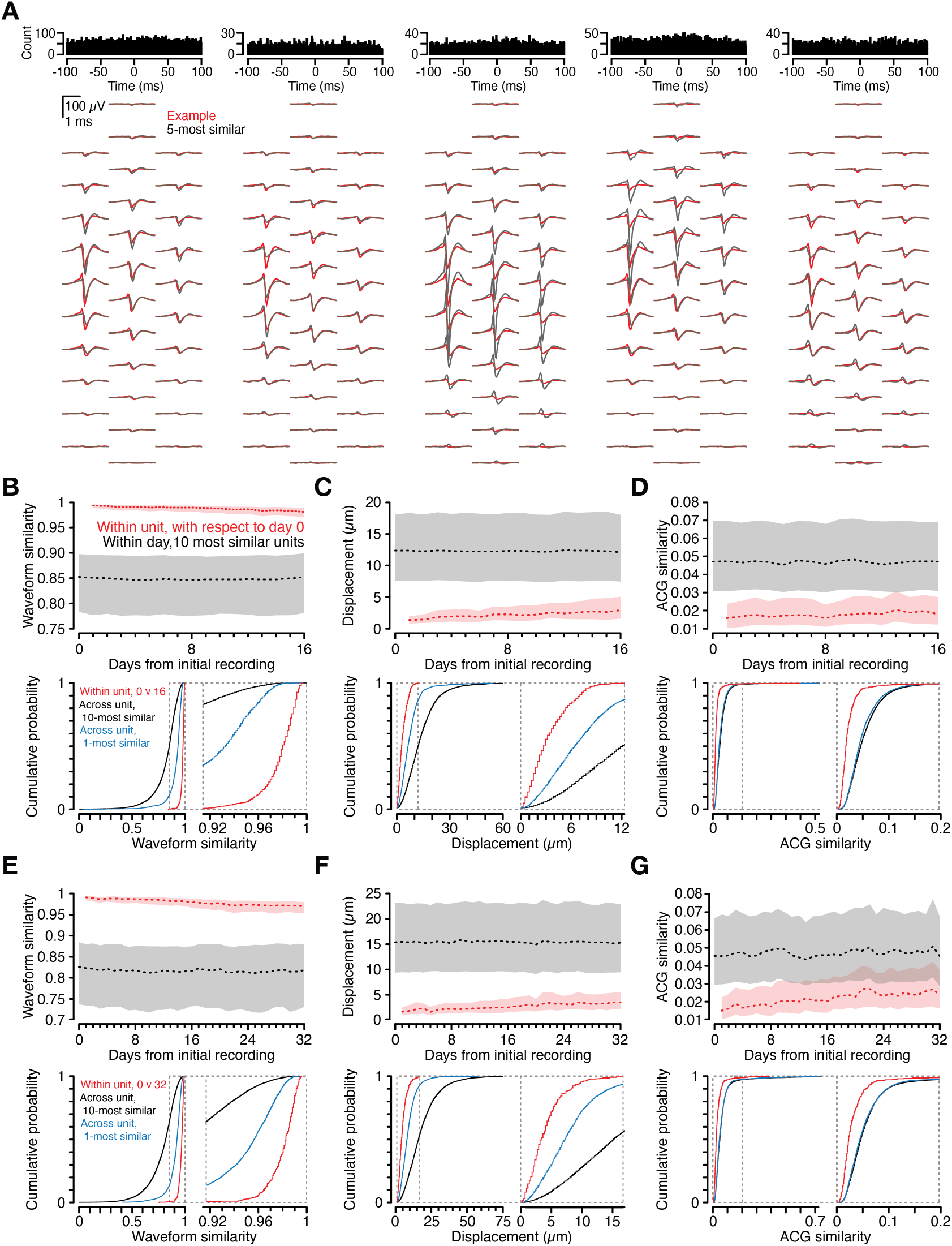
Single units are more similar to themselves across days than to any other single unit on the probe. **A**, Representative example single unit (red, same as shown in Fig. 1D) and the five most similar single units recorded on day 32 (black), as measured by the Pearson’s correlation between pairs of single unit waveforms. Top, spike time cross correlograms between the example single unit and each of the five most similar single units. The absence of a dip in cross correlogram amplitude at the 0 time lag (refractory period violations) indicates that the example single unit and each of the five most similar single units correspond to distinct neurons. **B**, Top, for single units held across days, Pearson’s correlation between waveforms (waveform similarity) measured on day 0 and waveforms measured on subsequent days (red), and waveform correlations between a given single unit and the ten most similar single units to it within a given day (black). Dashed line, median. Shading, indicates boundaries of top and bottom quartiles. Bottom, for single units held across days, cumulative distributions of waveform similarity for day 0 vs. day 16 (red), waveform similarity between a given single unit and those 10 most similar to it (black), and waveform similarity between a given single unit and the 1 most similar to it (blue). Median within-unit correlation between day 0 and day 16, 0.98 (Q1 = 0.97, Q3 = 0.99); median across-unit correlation with the 10 most similar, 0.85 (Q1 = 0.78, Q3 = 0.90); median across-unit correlation with the 1 most similar, 0.93 (Q1 = 0.90, Q3 = 0.95). **C** and **D**, Same analysis as **B** but for single unit centroid displacement and spike time autocorrelogram (ACG) similarity (Euclidean norm across ACG shape), respectively. **C**, Median within-unit displacement between day 0 and day 16, 2.9 μm (Q1 = 1.6 μm, Q3 = 5.0 μm); median across-unit distance from the 10 most similar, 12.3 μm (Q1 = 7.5 μm, Q3 = 18.0 μm); median across-unit distance from the 1 most similar, 6.5 μm (Q1 = 3.4 μm, Q3 = 9.4 μm). **D**, Median within-unit ACG similarity between day 0 and day 16, 0.018 (Q1 = 0.012, Q3 = 0.027); median across-unit similarity with the ACGs of the 10 most similar, 0.047 (Q1 = 0.031, Q3 = 0.069); median across-unit similarity with the ACGs of the 1 most similar, 0.044 (Q1 = 0.030, Q3 = 0.064). **E-G** same as **B-D** but for the experiments performed across a 32-day interval. **E**, waveform similarity; median within-unit similarity between day 0 and day 32, 0.97 (Q1 = 0.95, Q3 = 0.98); median across-unit correlation with the 10 most similar, 0.82 (Q1 = 0.73, Q3 = 0.88); median across-unit correlation with the 1 most similar, 0.92 (Q1 = 0.89, Q3 = 0.95). **F**, centroid displacement; median within-unit displacement between day 0 and day 32, 3.5 μm (Q1 = 2.1 μm, Q3 = 5.6 μm); median across-unit distance to 10 most similar, 15.4 μm (Q1 = 9.5 μm, Q3 = 22.9 μm); median across-unit distance to 1 most similar, 7.4 μm (Q1 = 4.5 μm, Q3 = 10.0 μm). **G**, ACG similarity; median within-unit ACG similarity between day 0 and day 32, 0.024 (Q1 = 0.016, Q3 = 0.038); median across-unit similarity with the ACGs of the 10 most similar, 0.047 (Q1 = 0.029, Q3 = 0.067); median similarity with the ACG of the 1 most similar, 0.045 (Q1 = 0.026, Q3 = 0.063). All within-unit metrics are significantly different from across-unit metrics (p < 5.0 × 10^−64^, Wilcoxon rank-sum), for both the 1-most and 10-most similar comparisons, across both the 16-day (N = 690 single units from 7 mice) and 32-day (N = 379 single units from 6 mice) interval experiments. Thus, single units are more similar to themselves across days than they are to even those single units most similar to them recorded within a given day.

**Extended Data Figure 3.**
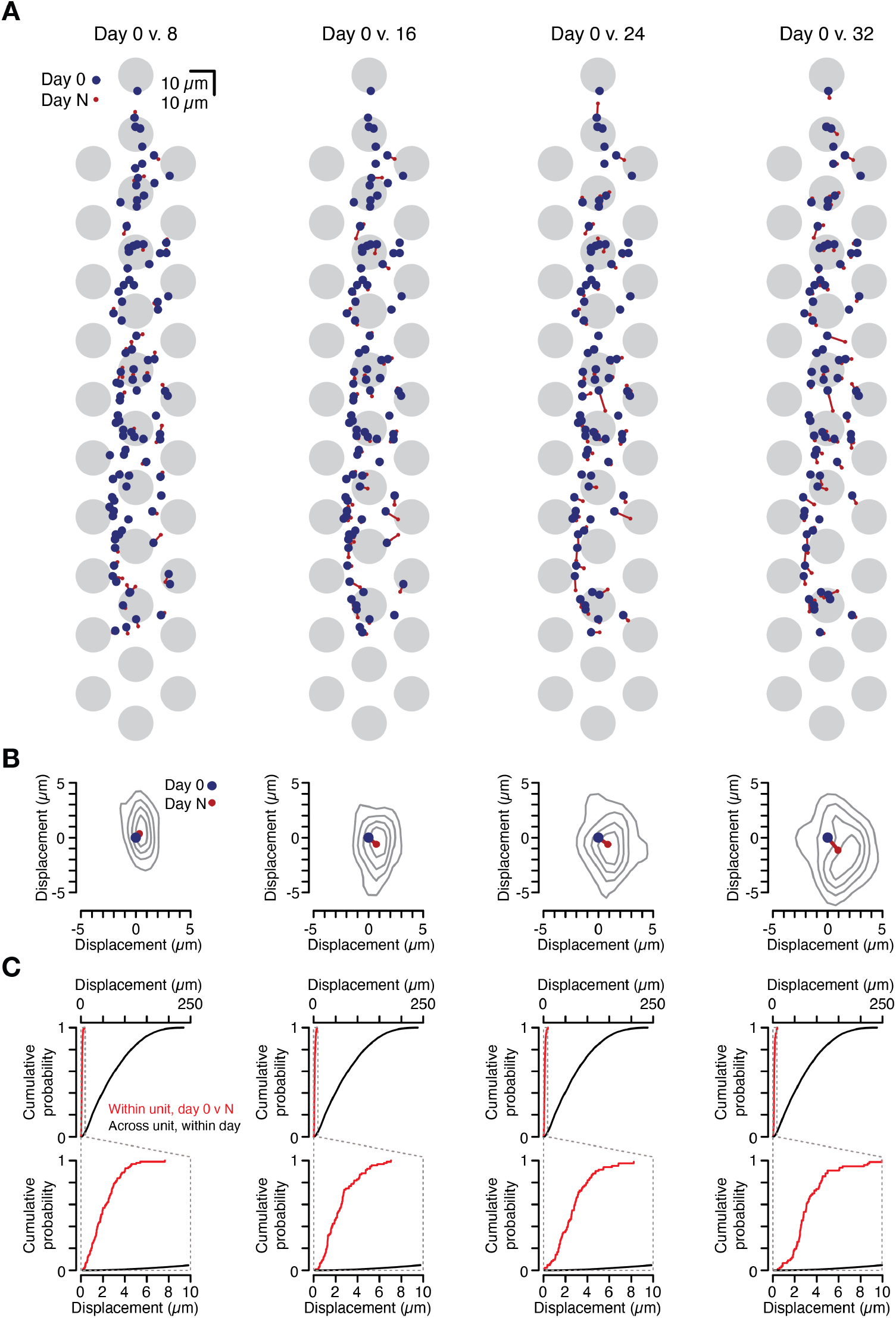
Estimated single unit position is stable across days. **A**, Single unit centroid positions across a 32-day interval from a representative animal. Centroid position for each of a population of single units measured on day 0 (blue circles) and days 8, 16, 24, and 32 (red circles, columns 1 through 4, respectively; days 0 vs. 8, N = 100 single units; days 0 vs. 16, N = 94 single units; days 0 vs. 24, N = 84 single units; days 0 vs. 32, N = 77 single units.) Single unit centroid is defined as the center of mass estimated from the amplitude of the mean waveform at each recording site. Grey circles indicate the size and position of the probe’s 32 electrodes. **B**, Mean displacement of single unit centroids from this animal between day 0 (blue circle, defined at origin) and days 8, 16, 24, and 32 (red circle, columns 1 through 4, respectively). Grey contours indicate quintile boundaries of the distribution of centroid position displacement between day N and day 0 for all single units in a given interval. **C**, Cumulative distribution of within-unit centroid displacement (red) between day 0 and days 8, 16, 24, and 32 (columns 1 through 4, respectively; days 0 vs. 8 within-unit median = 1.8 μm, across-unit median = 61.7 μm, p = 3.0 × 10^−66^; days 0 vs. 16 within-unit median = 2.1 μm, across-unit median = 62.8 μm, p = 4.3 × 10^−62^; days 0 vs. 24 within-unit median = 2.6 μm, across-unit median = 61.6 μm, p = 1.9 × 10^−55^; days 0 vs. 32 within-unit median = 2.8 μm, across-unit median = 61.7 μm, p = 8.6 × 10^−51^, Wilcoxon rank-sum) and across-unit centroid displacement within day (black) for this animal. Inset below shows both distributions for displacements between 0 μm and 10 μm.

**Extended Data Figure 4.**
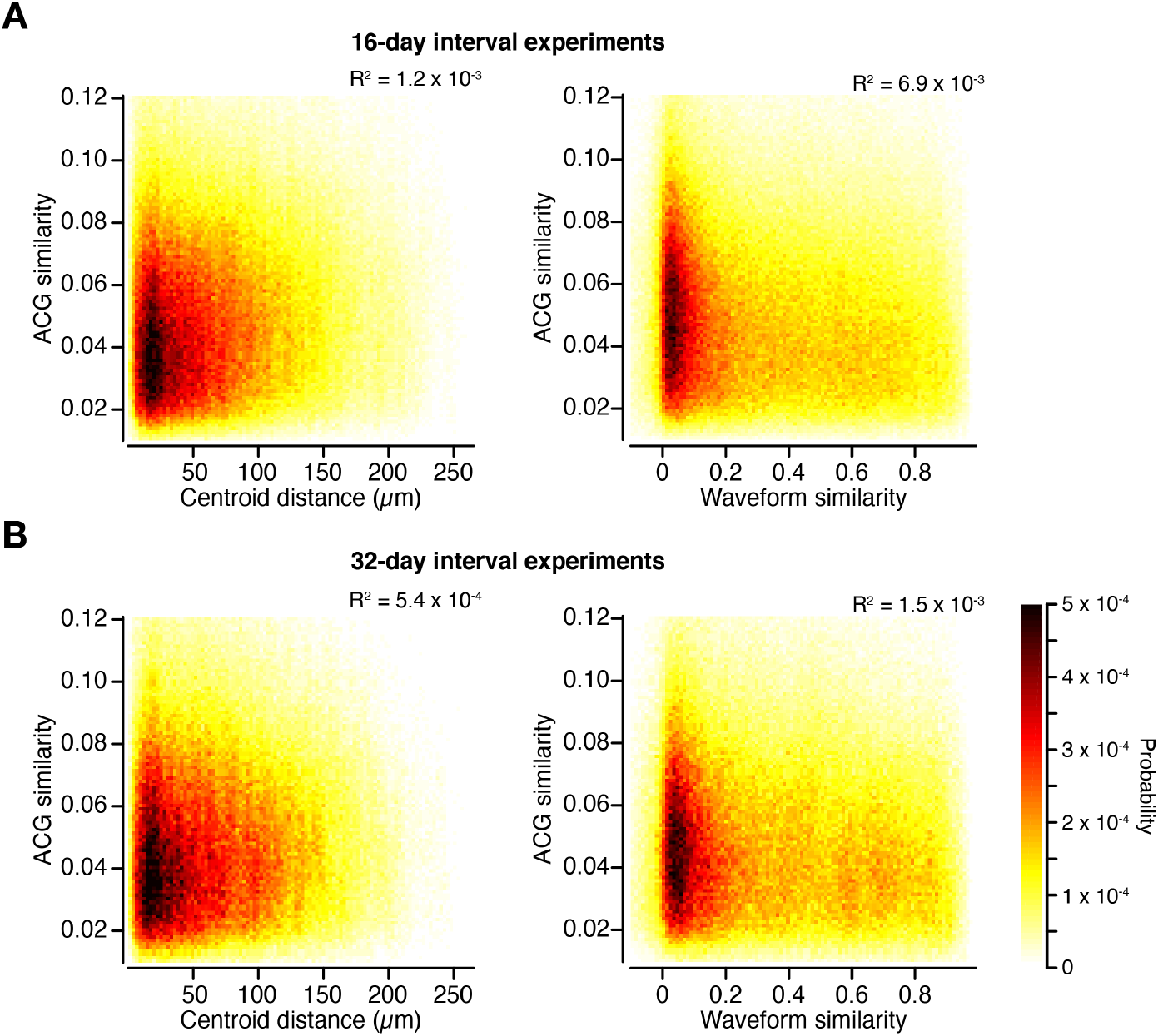
Waveform and non-waveform-based single unit features are uncorrelated. The similarity (Euclidean norm), on a given day, between the autocorrelograms (ACG) of a pair of simultaneously recorded single units vs. the similarity (Pearson’s correlation) between that pair’s spike waveforms (right) and the distance (Euclidian norm between the centroids) between the two single units (left), measured for **A**, 16-day interval experiments (N = 1,248,216 pairs of single units from 7 mice on 17 days) and **B**, 32-day interval experiments (N = 841,138 pairs of single units from 6 mice on 33 days). 3D histogram, with lighter colors indicating lower probability and darker colors indicating higher probability. This demonstrates that waveform-based features (waveform similarity and single unit centroid distance) vary independently of the similarity of the spike-time ACGs. Thus, ACG similarity is a measure of single unit stability independent of features to which the spike sorting algorithm is sensitive.

**Extended Data Figure 5.**
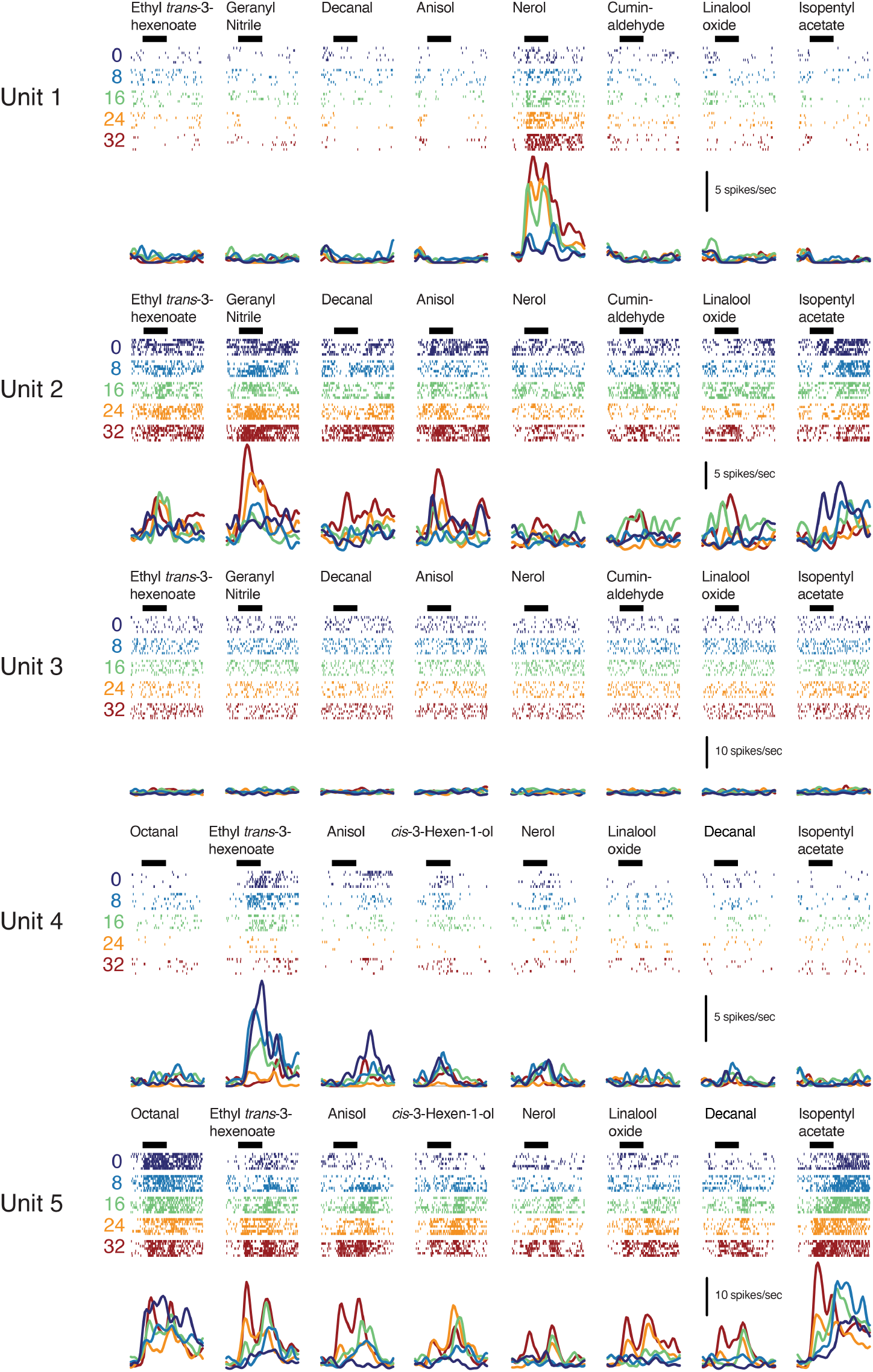
Evoked responses in single units across days. Responses of five single units across a 32-day interval selected to illustrate diversity of response profiles and drift rates. Columns separate test odorants (chemical names, top). Spike rasters (rows: 7 trials per day) and peristimulus time histograms are colored by day as indicated. Horizontal black bars, 4 sec odorant pulse.

**Extended Data Figure 6.**
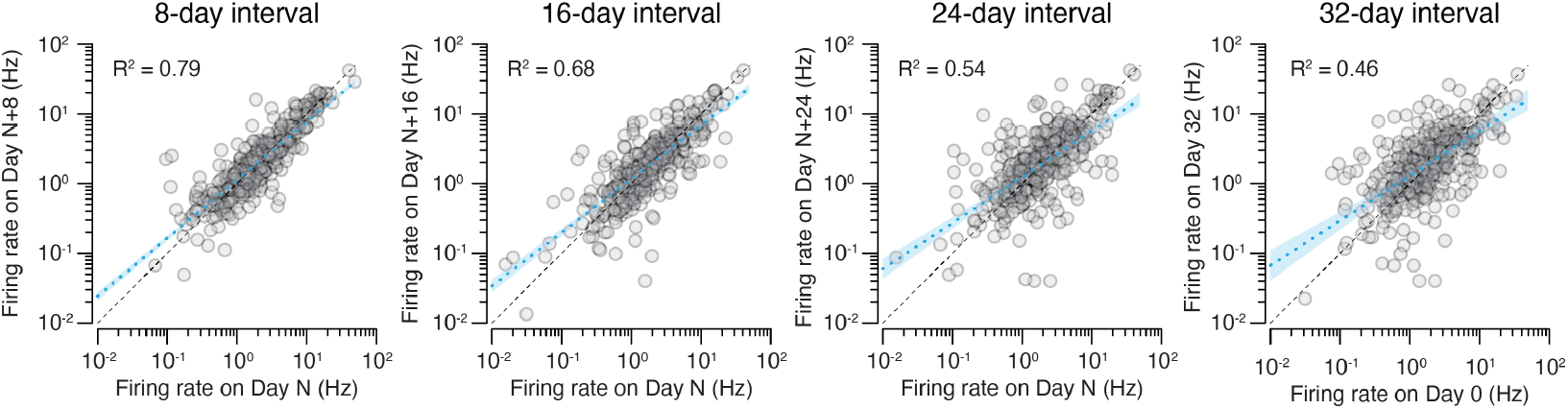
Spontaneous firing rates across days. Mean single unit spontaneous firing rate (baseline firing rate) across intervals of 8 days (*ρ* = 0.89, N = 2,177 single units), 16 days (*ρ* = 0.82, N = 1,412 single units), 24 days (*ρ* = 0.74, N = 816 single units) and 32 days (*ρ* = 0.68, N = 379 single units) from 6 mice. All correlations are significant (p < 4.0 × 10^−52^). Each plot shows a random subset of 379 single units, to match the number of single units recorded across the 32-day interval (right panel). Black dashed line, unity; blue dashed line, linear regression; blue shading, 95% C.I.

**Extended Data Figure 7.**
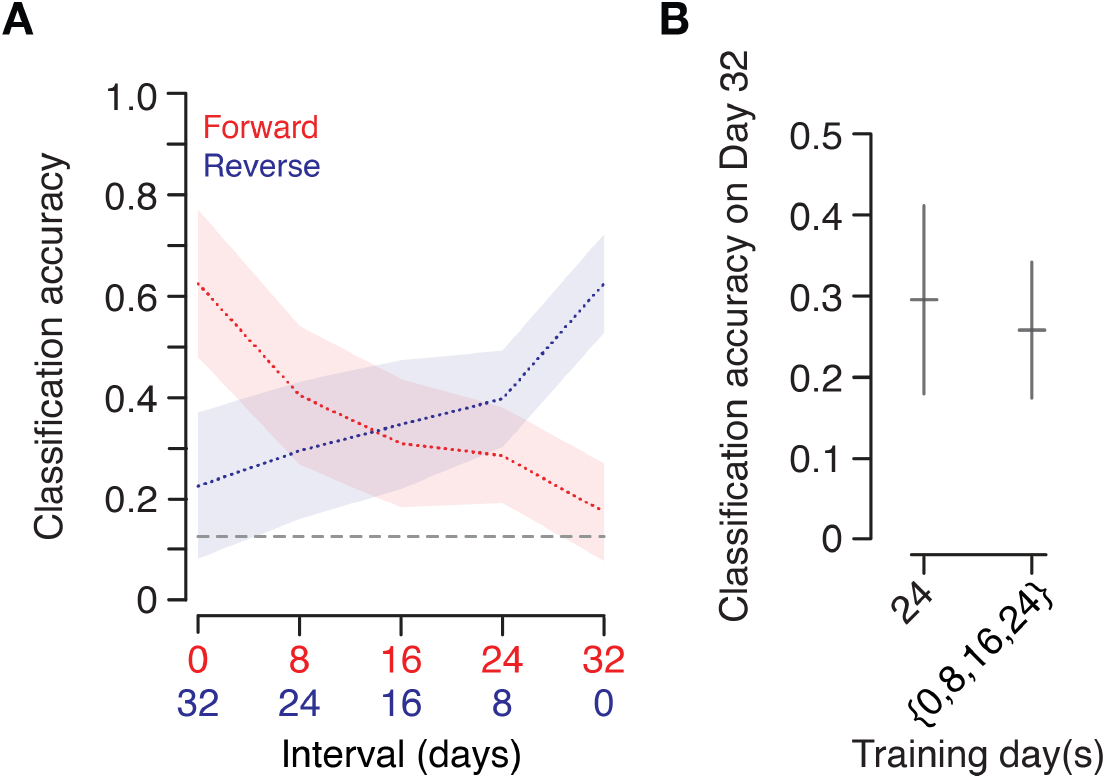
Drift is symmetric in time and linear classification across multiple training days does not improve performance. **A**, Classification accuracy (support vector machine, linear kernel) trained on earlier days and tested on later days (red, ‘forward’) compared with a model trained on later days and tested on earlier day (blue, ‘reverse’). Red and blue dashed lines, mean; Shading, standard deviation; Grey dashed line, chance performance; limit of 41 single units per animal with 100 permutations. **B**, Classification accuracy of a classifier trained on responses on day 24 alone (all 56 trials) and tested on day 32 compared with a model trained on 75 random subsets of 56 trials drawn from days 0, 8, 16, 24 and tested on day 32; 0.26 ± 0.08 multi-day accuracy vs. 0.30 ± 0.12 single-day accuracy, mean ± s.d., p = 2.6 × 10^−5^, Wilcoxon rank-sum, 100 random subsets of 23 single units per animal. For both **A** and **B**, N = 3 animals that were presented a test odorant panel of 8 stimuli. A support vector machine trained on concatenated data from days 0, 8, 16, and 24 will assign high weights to single units that were stable (less variable) across those four days and conversely assign low weights to single units whose responses varied. Thus, if there is a special population of neurons whose responses are informative about stimulus class and are more stable than others, a model trained on a concatenation of days 0, 8, 16, and 24 ought to perform better when tested on day 32 than a model trained on day 24 alone. However, we observe the opposite: it is not possible to establish single units most informative about odor identity on day 32 based on their responses across days 0 through 24. This argues against the presence of an informative stable subpopulation.

**Extended Data Figure 8.**
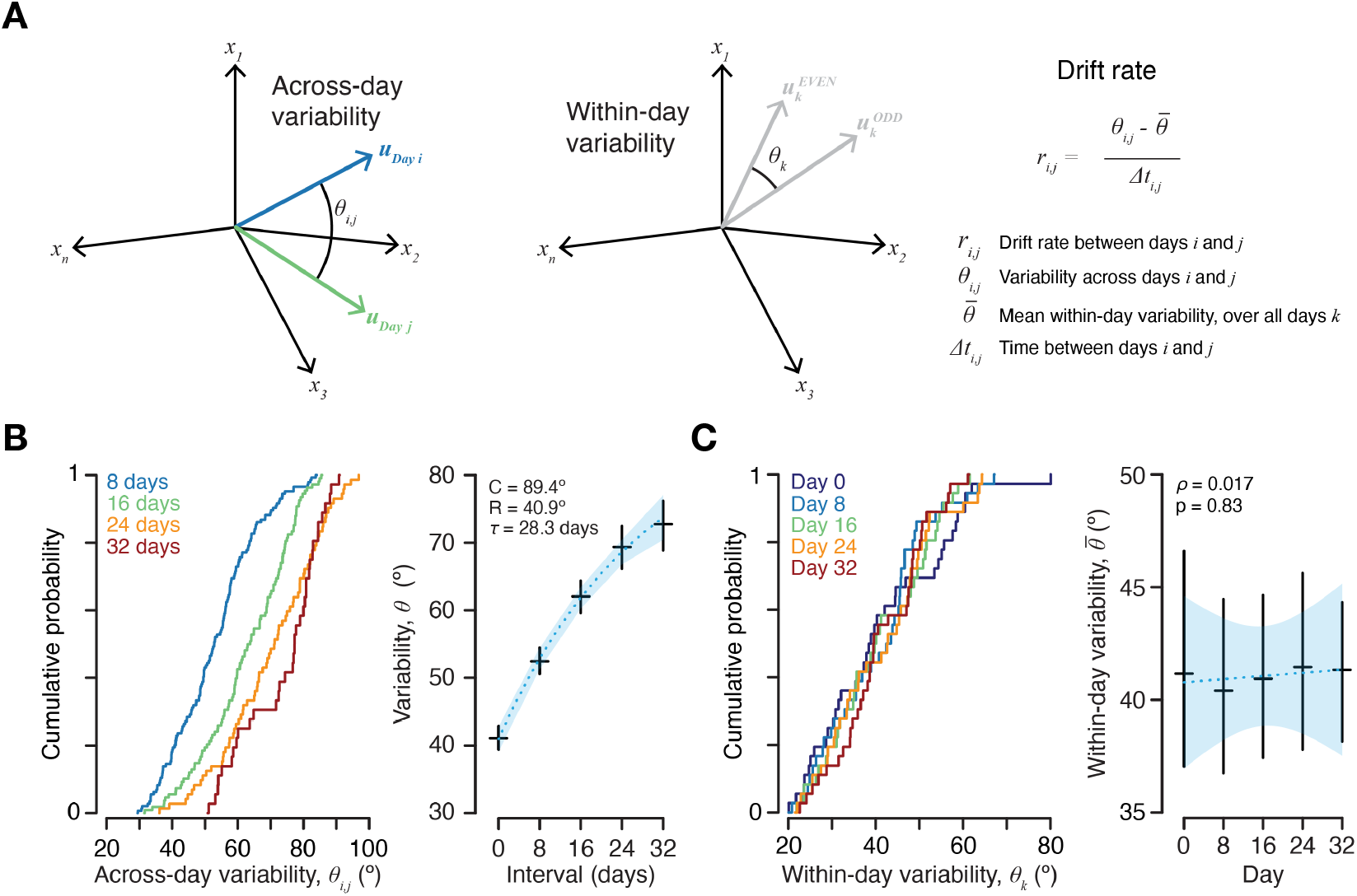
Estimating drift rate. Representational drift between any pair of days can be estimated by measuring the difference in representations across days and compensating for within-day variability^33^. **A**, Left, variability across days (across-day drift + within-day variability), estimated by computing the angle (*θ_i,j_*) between trial-averaged population vectors ***u***_*i*_ and ***u***_*j*_ for each odor across each pair of days *i* and *j*. Middle, variability within a day (noise), estimated by measuring the mean over all days of the angle between the trial-averaged population vectors for each odor within each day on odd trials versus even trials (*θ_k_*). Right, the drift rate is defined as the difference between the angle measured across days (*θ_i,j_*) and the mean angle measured within days, between even and odd trials, across all days *k* (*θ_k_*) divided by the time elapsed between days *i* and *j* (*Δt_i,j_*). This yields the drift rate (*r_i,j_*) expressed in degrees per day. **B**, Left, cumulative distribution of angles measured between trial-averaged population vectors across intervals of 8, 16, 24, and 32 days. Right, black crosses, population vector angle as a function of interval (mean across all odors ± 95% C.I., 0-day interval (within-day, odd vs. even trials), N = 180 pairs; 8-day interval, N = 144 pairs; 16-day interval, N = 108 pairs; 24-day interval, N = 72 pairs; 32-day interval, N = 36 pairs, all from 6 mice.) Blue dashed line, exponential regression curve, 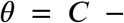 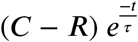, where *θ* is the variability (angle), *C* the asymptote, *R* the intercept at *t* = 0 (within-day variability), and *τ* the time constant of the exponential in days. Blue shading, 95% C.I. **C**, Left, cumulative distribution of within-day angles between trial-averaged population vectors for odd versus even trials for each day. Right, black crosses, within-day population vector angle for each day (mean across all odors ± 95% C.I., for each day N = 72 pairs from 6 mice). Blue dashed line, linear regression. Blue shading, 95% C.I. No pair of within-day angles differs significantly (p ≥ 0.56 for all pairs, Wilcoxon rank-sum).

**Extended Data Figure 9.**
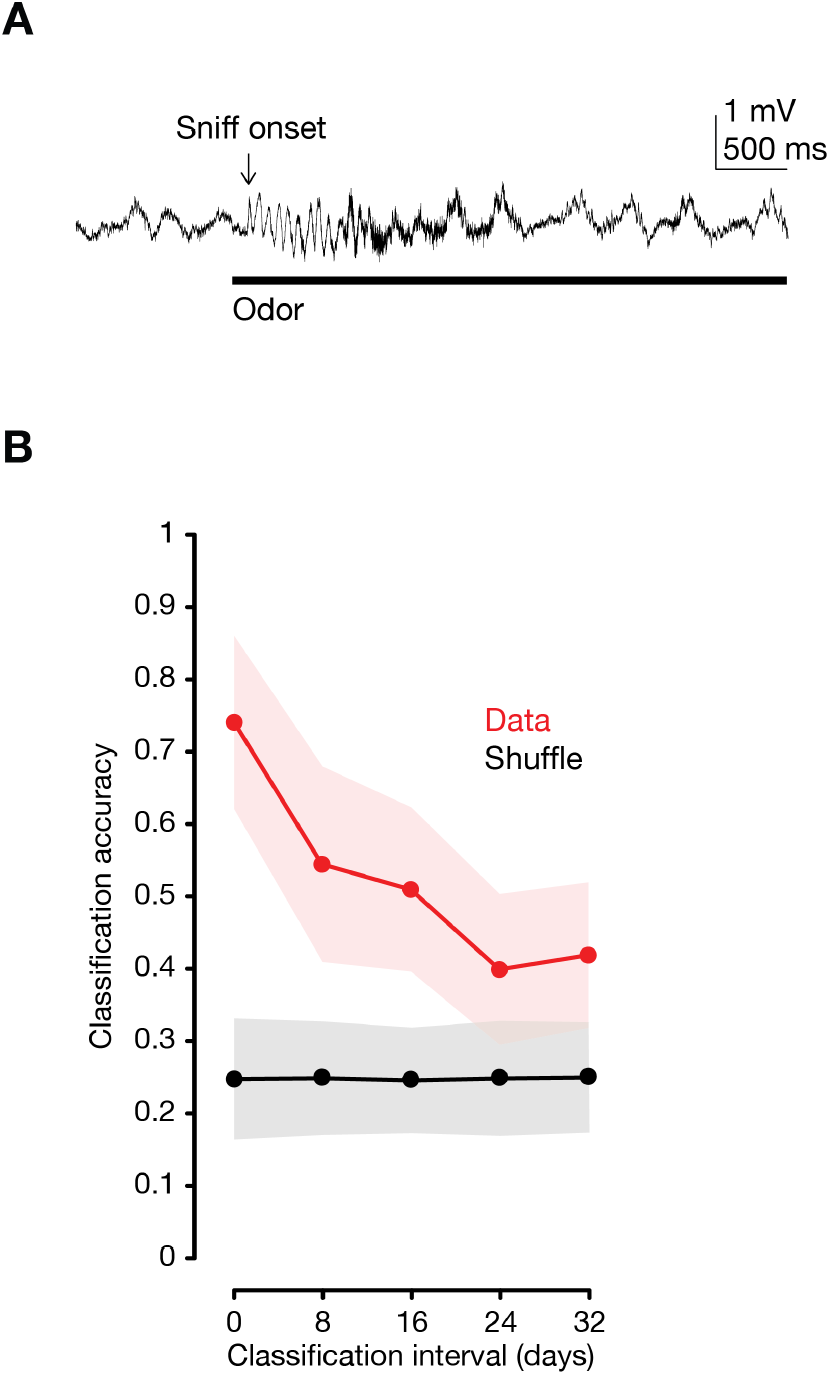
Activity during the early phase of the odor response drifts across days. **A**, Mean local field potential over all 32 electrode sites (filtered 0.1 – 20 Hz) during odorant administration. This oscillation was used on each trial to estimate the time of sniff onset. Only spikes occurring within the 190 ms following and 10 ms before the detection of sniff onset were analyzed. **B**, Red, classification accuracy (support vector machine, linear kernel) trained on day 0 and tested on each subsequent day for the first 200 ms of the odor response (N = 71 single units from 1 mouse, 4-way classification). Classification performance on day 0 computed using leave-one-out cross-validation. For all other days the model is trained on day 0 and tested on responses from the later day using *z*-scored responses. Points, mean; Shading, standard deviation.

## Methods

### Ethical compliance

All procedures were approved by the Columbia University Institutional Animal Care and Use Committee (protocol AC-AAAT5466) and were performed in compliance with the ethical regulations of Columbia University as well as the Guide for Animal Care and Use of Laboratory Animals.

### Stereotactic targeting and head plate attachment surgery

10-17 week old (12 ± 2 weeks, mean ± s.d., N = 45 mice) male C57BL/6J mice (Jackson laboratories, Bar Harbor, ME) were anesthetized with isoflurane (3% induction, 1.5-2% maintenance), placed on a feedback-controlled heating pad (Fine Science Tools, Foster City, CA) and secured within a stereotactic frame (Model 1900-B Head Holder Assembly with Sagittal Plate from a Model 1900 Stereotaxic Alignment Instrument, David Kopf Instruments, Tujunga, CA) affixed to an air table (TMC, Peabody, MA). All stereotactic targeting was performed using a separate motorized manipulator (Scientifica, Uckfield, UK) mounted on the air table under a Leica M80 dissection stereo microscope (Leica Microsystems, Wetzlar, Germany), with images captured using the Leica Application Suite. Carprofen (5 mg/kg) was administered via subcutaneous injection as a preoperative analgesic and bupivacaine (2 mg/kg) was delivered underneath the scalp to numb the area of the incision. The skull was exposed and cleaned with sterile cotton swabs.

To reliably target the anterior piriform cortex (APCx) we employed a series of stereotactic injection procedures to reduce targeting variability across animals. The skull was aligned by first adjusting the angle of the head, dorsal tilt (pitch), such that lambda and the rostral confluence of the sinus (RCS) both lay in the horizontal plane of the motorized manipulator. The position of the lateral edges of the frontal and parietal bones was mapped at 500 μm increments from 1,000 μm posterior to RCS (pRCS) to 4,500 μm pRCS. These measurements were compared to mean values and the angle of the head, coronal tilt (roll) and sagittal tilt (yaw), was adjusted using the head-holder assembly to minimize differences from a mean reference animal (N = 164 mice). The midline was defined as the midpoint between the lateral edges of the frontal bones at 1,000 μm pRCS, 1,500 μm pRCS, and 2,000 μm pRCS. If the difference between any of these landmarks exceeded two standard deviations from mean values, animals were not used for probe implantation.

Following head alignment, the area surrounding the target location for APCx was marked with hatch marks scored with a scalpel and then labeled with oil-based ink. The skull was covered in a thin layer of cyanoacrylate adhesive (Krazy Glue, Elmer’s Products, Atlanta, GA). A glass capillary was then positioned at the target for APCx (2,250 μm lateral to the midline, 1,150 μm pRCS) and a photograph was taken of the capillary position for positioning the silicon probe on the day of implantation using a camera attached to the dissection microscope.

A portion of skin adjacent the incision was removed and the area of skin and muscle surrounding the skull was temporarily covered with silicone elastomer (Kwik-Cast, World Precision Instruments, Sarasota, FL) to protect it during application of cement. A coating of adhesive luting cement (C&B-Metabond, Parkell, Inc., Edgewood, NY) was applied atop the layer of cyanoacrylate adhesive, covering the skull except for the probe target area. A titanium head plate (27.4 mm × 9.0 mm × 0.8 mm, G. Johnson, Columbia University) was lowered onto the skull using the micromanipulator. A custom adapter (G. Johnson, Columbia University) positioned the head plate within the horizontal plane of the micromanipulator. The headplate was secured with additional applications of luting cement, again ensuring that no luting cement covered the probe target area. A well was then formed using dental acrylic (Tru-Pak Orthodontic Acrylic, Stoelting, Wood Dale, IL) such that the entire edge of the opening of the headplate was in contact with cement and the probe insertion site was within this well, separated from any exposed tissue.

Two bone screws, each connected to 15 mm long 36 AWG wire (Phoenix Wire, South Hero, VT) soldered to an Amphenol pin (A-M Systems, Carlsborg, WA) trimmed to 1 mm, were then inserted bilaterally above the cerebellum opposite the midline to serve as ground and reference electrodes. The bone screws were then sealed in place with luting cement and the wires were directed to the anterior portion of the headplate, opposite the probe target area, and sealed in position with dental acrylic. The Kwik-Cast was removed and the incision posterior to the headplate was closed with sutures. Mice were allowed 6 - 35 days (19 ± 10 days, mean ± s.d., N = 45 mice,) to recover before attempting probe implantation.

### Probe preparation

All recordings were performed using A1×32-Poly3-5mm-25s-177 silicon probes (177 μm^2^ site surface area, 3-column honeycomb site geometry with 18 μm lateral and 25 μm vertical site spacing, 36 μm center-to-center horizontal span, 275 μm center-to-center vertical span, 114 μm maximum shank width near the sites, 15 μm shank thickness) with an H32 connector (NeuroNexus Technologies, Ann Arbor, MI). Prior to implantation a NanoZ (White Matter, LLC, Seattle, WA) was used to electroplate each of the 32 electrodes (+5 nAmp, 30 sec) with poly(3,4-ethylenedioxythiophene) (PEDOT) to reduce site impedence^56,57^.

The probe tip base was affixed with cyanoacrylate adhesive to a 3D-printed titanium anchor, which itself was coupled to a probe tip connector, 3D-printed using a water-soluble polymer (polyvinyl alcohol, PVA). The probe tip connector was attached to custom-designed fittings (T. Tabachnik, Columbia University), which also held the probe’s printed circuit board (PCB) and Omnetics connecter and were secured to a 2-axis gimbal (G. Johnson, Columbia University).

The 2-axis gimbal was mounted on the Scientifica manipulator and was employed to align the probe shank to the vertical axis of travel of the manipulator. Alignment was complete when the probe shank, as it passed through the approximately 3 mm that would enter the brain during implantation, deviated no greater than 10 μm from a fixed point, visualized using a reticle (alignment within 12 arcminutes). The probe was then left, mounted, on the 2-axis gimbal with the alignment preserved. During the implantation procedure, the 2-axis gimbal was again mounted on the Scientifica manipulator and the probe’s alignment was confirmed before initiating the penetration.

### Probe implantation

Mice were anesthetized with ketamine/xylazine (100 mg/kg ketamine and 10 mg/kg xylazine initial dose). An intraperitoneal cannula was inserted to administer maintenance doses of ketamine (30 mg/kg every 20-30 mn). The animal was placed on the feedback-controlled heating pad and carprofen (5 mg/kg) was administered via subcutaneous injection. The headplate was then inserted in the stereotactic frame using custom holders and leveled to ensure its alignment to the horizontal plane of the Scientifica manipulator. The alignment of the headplate was confirmed using a dial test indicator affixed to the Scientifica manipulator. To minimize vibration during implantation all procedures were performed on an air table (TMC, Peabody, MA).

Using the photographs previously taken during head-plate attachment, a glass guide pipette attached to the manipulator was positioned at the previously targeted location for probe implantation. A craniotomy was then performed using a dental drill (Osada Success 40, Osada Electric Co., Ltd., Tokyo, Japan) and a 0.5 mm drill bit (Fine Science Tools, Foster City, CA). Considerable care was taken when performing the craniotomy to avoid damage to the underlying tissue; in the event of pial bleeding, edema, or other signs of damage to the brain the probe implantation was aborted. After successfully performing the craniotomy, the glass guide pipette was lowered to the surface of the brain and its position relative to the blood vessels on the pial surface was photographed to guide later placement of the silicon probe. The craniotomy was then covered with cotton moistened with phosphate buffered saline. A custom-designed well molded out of silicone elastomer (Kwik-Cast) was affixed atop the head plate to permit later dissolution of the PVA probe tip connector once the probe was cemented in place. The glass guide pipette was then removed from the manipulator and the silicon probe was positioned above the insertion site using the photographs of the guide pipette.

The probe was lowered 1,000 μm below the pial surface under manual control of the motorized manipulator. Throughout the penetration neural signals were monitored, both visually and aurally. Beyond depths of 1,000 μm below the pial surface, software was used to control the speed of descent. The probe was lowered from 1,000 μm to 2,200 μm at 2 μm/sec and beyond 2,200 μm was advanced at 1 μm/sec until the desired target area was reached. Proper probe alignment was confirmed by visually tracking the movement of distinctive spike waveforms as the array of sites passed by them during penetration. Good alignment was further confirmed by temporarily retracting the probe (between approximately 500 μm and 1500 μm) and confirming that spike waveforms that the probe sites had already passed reappeared. In case signals were attenuated during these two checks, we concluded that the probe was causing damage to the tissue, possibly due to slight misalignment. Implantation was then aborted.

The cell dense layer 2 of APCx was identified by a rapid increase in threshold crossing events and tight coupling of the local field potential to the animal’s breathing cycle. In 15 preliminary APCx targeting experiments we coated a glass capillary with DiI (Thermo Fisher Scientific, Waltham, MA) and determined post-hoc in histology appropriate stereotactic coordinates. Then, in 5 preliminary acute mapping experiments, during which we applied DiI to the back of the probe, we related clear physiological signatures along the penetration track with physical locations determined post-hoc in histology (Fig. 1A). These physiological signatures permit us to determine unambiguously whether our electrode sites are located in the cell-dense layer of piriform, and thus to decide whether to commit a probe to chronic implantation. Once the probe reached its target position (depth from pial surface 3,118 ± 131 μm, N = 32 mice, mean ± s.d.) it was retracted to accommodate settling of the tissue (retraction from target depth 117 ± 41 μm, mean ± s.d., N = 32 mice). If the probe failed to target the cell dense layer 2 on the first attempt, implantation was aborted; to minimize tissue damage, we never made more than a single penetration during implantation.

A thin protective coating of uncured (liquid) silicone (DOWSIL 3-4680 Silicone Gel, Dow Inc., Midland, MI) was then carefully applied to the pial surface via a 23-gauge blunt syringe needle affixed to a 1 ml syringe. After the silicone had cured, very liquid (uncured) dental acrylic (Tru-Pak Orthodontic Acrylic, Stoelting, Wood Dale, IL) was carefully applied via a 23-gauge blunt syringe needle affixed to a 1 ml syringe, surrounding the shank of the probe up to its base. The dental acrylic was then allowed to cure for at least 20 mn. The dental acrylic and still exposed probe tip holder were then carefully covered in Grip Cement (Dentsply, York, PA), again via a 23-gauge blunt syringe needle affixed to a 1 ml syringe, in two phases. The Grip Cement was allowed to harden for at least 20 mn after each phase. The Grip Cement surrounded the probe tip up to the PVA probe tip connector and flexible polyamide cable that connected the probe tip to the probe’s PCB and Omnetics connector. Between the time the probe entered the brain and the time it was gently decoupled from the micromanipulator (see below) considerable care was taken to minimize movement of the probe relative to the brain tissue in any dimension other than vertical. Only fluids (uncured silicone, dental acrylic, Grip Cement) were permitted to contact the probe/micromanipulator assembly; contact with a solid object (e.g. 23-gauge needle) typically resulted in degradation of the continuously monitored spike waveforms and abortion of the probe implantation.

The silicone well was then filled with distilled water in order to dissolve the PVA probe tip connector, thus ensuring gentle decoupling of the probe from the micromanipulator. Once the PVA had dissolved (~30 mn), the only remaining physical connection between the probe tip and the PCB and Omnetics connector was the thin, flexible polyimide cable. Thus, at this point, the PCB and Omnetics connector could be disconnected from the manipulator without disturbing the implant. The PCB and Omnetics connector were then gently affixed with Grip Cement within a copper mesh casing, itself affixed to the headplate using Grip Cement. Importantly, the PCB and Omnetics connector were not affixed to the headplate or Grip Cement surrounding the probe itself, to isolate the implant from physical insult when plugging/unplugging the headstage during subsequent recording. The copper mesh casing was then secured to the headplate using Grip Cement and a lead connected to the probe’s ground wire was soldered to the copper mesh. The probe was unplugged from the headstage and to protect the pin contacts from debris and dander the exposed sites of the Omnetics connector were covered with tape (Scotch Magic Tape, 810-NA, 3M, Saint Paul, MN). The animal was then returned to its homecage.

### Stabilization following implantation

We have observed that in some cases the brain tissue appears to relax around the probe over as long as 2-3 weeks, as evidenced by large, distinctive single unit waveforms drifting up the vertical axis of the electrode sites. In order to ensure complete stabilization of the tissue around the probe and thus permit longitudinal recordings of single units across days, mice were given 9 ± 3 weeks (mean ± s.d., minimum 5 weeks, N = 16 mice) to recover (Extended Data Fig. 1A, E). Thereafter, we began daily recordings. During an initial monitoring phase, animals were head fixed and the head stage was plugged into the probe’s Omnetics connector for daily 30 mn recording sessions to assess single unit stability across days. After at least 10 days of daily monitoring, experiments commenced (see Experiment schedule).

### Histology

Animals were transcardially perfused with 4% paraformaldehyde 0.1 M phosphate buffer fixative. The brain was postfixed overnight, washed in phosphate buffered saline and sectioned along the axial plane (100 μm sections) using a vibratome (Leica VT1000s, Leica Microsystems, Wetzlar, Germany). Cell bodies were stained using NeuroTrace (Life Technologies, Carlsbad, CA) for 30 mn at a 1:50 dilution. Sections were imaged under a Zeiss LSM 510 microscope (Carl Zeiss, Oberkochen, Germany) under the control of ZEN imaging software (Carl Zeiss).

### Experimental apparatus

During all recording sessions, animals were head fixed and positioned atop a trimmed absorbent underpad (Fisher Scientific, Hampton, NH) within a 3D printed polylactic acid tube (45.5 mm inner diameter, 49 mm outer diameter, 7 cm long, with a sealed back). For most experiments the tube was secured in a fixed position with respect to the headplate holders but to monitor behavioral responses to aversively conditioned stimuli the tube floated on a pair of frictionless air bearings using a Virtual Burrow Assay^37^. A background of bandpass filtered acoustic white noise (1000–45000 Hz; approximately 7 dB) was played throughout all experiments and the animal and apparatus were within a custom made sound attenuating chamber resting on an air table (TMC, Peabody, MA). Experiments were conducted under conditions of visual darkness, with animals illuminated with infrared light to permit simultaneous video monitoring.

### Stimulus delivery

Odorant stimuli were administered using a 4 sec odorant pulse with a 60 ± 20 sec inter-trial interval (ITI). Each odor test session consisted of 7 stimulus presentation blocks in which each odorant stimulus was delivered pseudorandomly within each block. Animals were given 3 mn to acclimate to head fixation before we launched stimulus presentation.

Odorant stimuli were administered using a custom built olfactometer. At all times a constant flow of air (0.7 l/mn) was directed through a nose port constructed of polyether ether ketone (PEEK), placed approximately 1 mm away from the nose of the animal. Air was routed to the nose port via one of two independent air lines: an air stream, normally routed to the nose port, and an odor stream, normally routed to exhaust but routed to the nose port upon administration of odorant stimuli. Both the odor and air streams were supplied with dedicated tanks of medical grade air and the flow rate of each was regulated by a dedicated mass flow controller (GFCS-010201, Aalborg, Orangeburg, New York). Both lines were directed through 50 ml glass bottles containing 15 ml dipropylene glycol (DPG, MilliporeSigma, Burlington, MA).

Monomolecular odorants were dissolved in 15 ml DPG in separate 50 ml bottles and replenished before each odor test session. The inlet and outlet of all odorant bottles were connected to the odor stream via two PEEK manifolds (Western Analytical, Vernon Hills, IL). The inlet of each bottle was affixed with a normally closed two-way valve. The outlet was affixed with a three-way valve normally open to an exhaust line (two-way and three-way valves manufactured by the Lee Company, Westbrook, CT). In preparation for odorant stimulus delivery, the two-way inlet valve was open and the three-way outlet valve was switched from the exhaust to the odor stream. The odor stream was then allowed to stabilize for at least 30 sec before administering the odorant stimulus. To deliver odorant stimuli, a four-way valve (NResearch Inc., West Caldwell, NJ) routed the air stream to exhaust, replacing it with the now odorized odor stream; the odorant stimulus was switched off when the four-way valve routed the odor stream back to exhaust and the air stream back to the nose port. The four-way valve was housed outside the experiment chamber in a sound attenuating box. The timing of olfactometer valve state changes and the trial structure of the experiment were controlled by a National Instruments board (National Instruments, Austin, TX) using software written in Python (PyDAQmx^58^ was used to interface with the NI-DAQmx driver).

After passing through the nose port all gas was routed to a photoionization detector (miniPID, Aurora Scientific, Aurora, ON, Canada). To minimize contamination, all material in contact with the odorized air stream, including the wetted material of all valves, was constructed of either Teflon, Tefzel, or PEEK. The flow of the air and odor streams were equalized before each experiment (using mass flow meter GFMS-010786 from Aalborg, Orangeburg, NY) and the tubing carrying the two streams from the four way valve was set to equal length and impedance to minimize variation in flow rate upon switching between the air and odor streams.

### Odorant stimuli

The following odorants were employed in these experiments, with their concentration (volume by volume in DPG) titrated during preliminary concentration dependence experiments to evoke responses in comparable fractions of piriform neurons: 2% *cis*-3-hexen-1-ol, 2% octanal, 2% anisole, 4% ethyl *trans*-3-hexenoate, 2% or 4% isopentyl acetate, 4% nerol, 4% decanal, 4% or 6% linalool oxide, 10% methyl salicylate, 10% cuminaldehyde, 10% geranyl nitrile (1:1 mixture of (E)- and (Z)-isomers), 10% R-(−)-carvone. All odorants were purchased from MilliporeSigma, Burlington, MA with the exceptions of nerol, ethyl *trans*-3-hexenoate, and cuminaldehyde, which were purchased from Thermo Fisher Scientific, Waltham, MA and geranyl nitrile, which was purchased from W. W. Grainger, Lake Forest, IL and manufactured by Tokyo Chemical Industry, Tokyo, Japan.

For measurements of representational drift over a 32-day interval, we tested two cohorts of mice. One cohort (N = 3 mice) was presented a panel of eight odorants every eight days. The second cohort (N = 3 mice) was presented a panel of four odorants every eight days, as well as a panel of four familiar odorants presented daily from days 0 to 16 and at 8-day intervals thereafter. For this second cohort only responses to the four odorants presented at 8-day intervals throughout were considered in our analysis of representational drift over a 32-day interval (Figs. 1–3). In one animal from the first cohort, the panel of eight odorants was presented once additionally, nine days before odor testing commenced. Otherwise, all odorants included in our measure of drift over a 32-day interval were not presented to the animals before the first test day (day 0).

For experiments measuring the effect of aversive conditioning on drift rate (Fig. 4A-C) the CS+ was nerol; the CS− was linalool oxide; the neutral odorants were isopentyl acetate, methyl salicylate, cuminaldehyde, and R-(−)-carvone. For experiments measuring the effect of daily experience on drift rate (Fig. 4D, E) the four familiar odorants included: *cis*-3-hexen-1-ol, octanal, anisole, ethyl *trans*-3-hexenoate, decanal, and geranyl nitrile (1:1 mixture of (E)- and (Z)-isomers). The four unfamiliar odorants included: isopentyl acetate, nerol, decanal, linalool oxide, methyl salicylate, cuminaldehyde, geranyl nitrile (1:1 mixture of (E)- and (Z)-isomers), and R-(−)-carvone. In all cases the animal had not previously experienced the unfamiliar odorant stimuli when the experiment commenced.

We avoided employing odorant stimuli that had been previously shown to elicit innate attraction or aversion and selected a structurally diverse array of odorant molecules with distinct organoleptic properties.

### Experiment schedule

For experiments monitoring representational drift across a 32-day interval, test odorants were administered every eight days, presented on five ‘odor-test days’, (days 0, 8, 16, 24, and 32). On intervening days, animals were plugged into the recording apparatus under identical conditions as on odor-test days but no odorant stimuli were administered. Except when otherwise noted, we grouped analysis between days 0-8, 8-16, 16-24, and 24-32 (8-day intervals) days 0-16, 8-24, and 16-32 (16-day intervals), and days 0-24, days 8-32 (24-day intervals).

For experiments monitoring the effect of daily exposure on representational drift, animals were presented a panel of four ‘familiar’ odorants daily across a 16-day interval (days -16 to 0). On day 0, in addition to the four ‘familiar’ odorants, a panel of four ‘unfamiliar’ odorants was presented that the animals had not previously experienced. For ‘cohort A’ (N = 5 mice), daily presentation of the ‘familiar’ odorants continued, with presentation of the ‘unfamiliar’ odorants occurring only every eight days (on days 0, 8, and 16). For ‘ cohort B’ (N = 5 mice) both ‘familiar’ and ‘unfamiliar’ odorants were presented every 8 days (days 0, 8, and 16). All analyses were performed on odor responses obtained on days 0, 8, and 16 (Extended Data Fig. 1A), looking at two 8-day intervals (days 0-8, days 8-16) and one 16-day interval (days 0-16).

### Odor-shock conditioning and testing

Conditioning and testing was performed using methods previously described^37^. Briefly, implanted mice were placed in a fear conditioning chamber under conditions of darkness with an acoustic background of white noise and allowed to acclimate for 5 mn. Eight blocks of CS+ and CS− odorants were presented in pairs of pseudorandomly interleaved trials. The odorant stimuli were 10 sec in duration with a 300 ± 100 sec ITI. During the final 2 sec of presentation of the CS+ only, the floor of the fear conditioning chamber was electrified (intensity 0.8 mAmp). Upon completion of all 8 trials, the mouse was permitted to recover for 5 mn in the fear conditioning chamber and then returned to its home cage.

One day after the fear conditioning protocol (day 0), as well as 16 days after (day 16), animals were placed within the Virtual Burrow Assay (VBA)^37^ to monitor their behavioral responses to odorant stimuli (CS+, CS−, and four neutral odorants) and record neural signals. Animals were acclimated to the VBA for 5 mn after head fixation (3 mn in the ‘open loop’ and 2 mn in the ‘closed loop’ configuration, see Ref. 37). The odorants were presented as in previously described experiments (4 sec odorant pulse, 60 ± 20 sec ITI). The position of the virtual burrow was measured using a laser displacement sensor. Each trial began with the animal resting outside the tube. Movement of the virtual burrow around the body during the odorant stimulus constituted ‘ingress’, akin to a flight response^37^. The position of the virtual burrow was quantified during the final second of odorant administration to measure the response to the CS+, CS−, and neutral odorants. On all intervening days (days 1-15) neural signals were recorded for at least 45 mn with the tube in a fixed position and without presentation of odorant stimuli.

### Data acquisition and spike sorting

Neural signals were acquired using a Cerebus Neural Signal Processor (Blackrock Microsystems, Salt Lake City, UT) using differential recordings with a CerePlex M headstage (bandwidth: 0.3 Hz – 7.5 kHz) sampled at 30 kHz under the control of the Cerebus Central Suite software. Continuous signals were then digitally bandpass filtered at 500 - 6000 Hz and preprocessed to subtract electrical artifacts associated with muscle activity in the jaw and/or mystacial pad: at each time bin the voltage registered on the seven electrode sites closest to 0 μV was averaged, and subtracted from all 32 electrode sites. Due to the magnitude of fields generated by piriform pyramidal neuron action potentials combined with the dense spacing between electrode sites, action potentials typically register on a large fraction, but not all, of the probe’s 32 electrodes. The choice of seven electrodes reflects a balance between accurate estimation of electrical artifacts common across all sites, and avoiding subtracting signals associated with veridical action potential events.

We then concatenated all recordings across all days included in a given analysis (e.g. for recordings of responses over a 32-day interval data from all 33 days of recording were concatenated in the order the recordings were acquired). The concatenated data was then spike sorted using the template matching algorithm Kilosort^59^ (github.com/cortex-lab/KiloSort); thus, the spike sorting algorithm was blind to recording day and treated data concatenated across days as a single continuous signal. We employed 256 templates with six passes and a spike threshold of 4*σ*. We did not employ any drift correction, therefore any drifting waveforms were assigned separate templates. The output of Kilosort was manually curated using Phy (github.com/cortex-lab/phy). During this manual curation phase we merged pairs of templates whose cross-correlogram indicated the spiking of a single common neuron; we did not perform any template splitting in the low dimensional visualization, instead relying on oversplitting by requesting such a large number of templates. We excluded from analysis any templates corresponding to electrical noise, as well as any templates for which refractory period violations exceeded 2%, (refractory period defined as an inter-spike interval <1.5 msec; median 0.08% refractory period violations, Q1=0.01%, Q3=0.24%). 17% of templates had zero refractory period violations. In sum, we did not attempt to stitch templates across multiple days, but rather designed the spike sorting pipeline to treat concatenated continuous signals as a single recording.

We then excluded from further analysis single units that were not stable across the recording. First we eliminated single units whose firing rate was near zero for any of the comparison days, as this indicates that the single unit was either gained or lost. Next we computed the correlation between each single unit’s waveforms measured on individual comparison days. For this, we computed each single unit’s mean waveforms (82 samples, 2.7 ms total duration), averaged on each day for each electrode, and then concatenated the mean waveforms from the probe’s 32 electrodes. Single unit waveform similarity is defined as the Pearson’s correlation between a pair of concatenated average waveform vectors. Within a given recording day, the 99th percentile correlation between the waveforms of different single units is 0.93. We therefore excluded single units whose correlation across a pair of comparison days fell below this value. Variations in waveform shape are amplified in very large amplitude single units. Thus, we employed a final manual check to identify and include those rare cases in which very large amplitude single units had across-day waveform correlations that fell slightly below 0.93. Each of these exceptions can be visualized in Extended Data Figure 1B, bottom, and 1F, bottom.

### Assessment of single unit stability across days

We employed three metrics to quantify the stability of single units across days: waveform similarity (computed as described above), displacement of the estimated single unit position relative to the probe sites, and similarity in the shape of the spike-time autocorrelogram (ACG).

To estimate single unit position, we computed a spatial average across electrode positions weighted by the mean waveform amplitude at each electrode. Many single units registered low amplitude waveforms on a majority of sites on the probe, which can confound the estimate of the centroid and cause underestimation of centroid displacement across days. For this reason we employed the square of the waveform peak-to-peak amplitude on each site to minimize the influence of low amplitude waveforms, estimating single unit centroid position (*x*,*y*) as

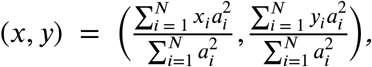

where *N* is the number of electrodes, *x_i_* is the lateral position of the *i*^th^ electrode, *y_i_* is the vertical position of the *i*^th^ electrode, and *a_i_* is the peak-to-peak amplitude of the spike waveform recorded at the *i*^th^ electrode. Single unit displacement was then computed using the Euclidean distance between single unit centroids on separate days. We note that due to the diverse and complex morphologies and biophysics of the neurons we recorded from, it is not possible to establish with precision the location of the source relative to the probe sites.

We computed the spike-time ACG over a 2 sec window using a bin size of 1 ms. To measure the similarity in the shape of the ACG we computed the Euclidean distance between normalized, filtered and then logarithmically binned spike-time ACGs.

For a given pair of comparison days, we computed each of these three metrics for all single units that were stable across that interval, as well as across all pairs of single units within a given recording session. We then compared within-unit/across-day distributions to the across-unit/within-day distributions to assess whether single units were more similar to themselves across days than to other single units simultaneously recorded within a day (Fig. 1E, Extended Data Figs. 1–3). We also identified for each single unit the ten, and one, most similar simultaneously recorded single unit(s) within a given day by ranking the waveform similarity for each single unit relative to every other single unit recorded on that day (Extended Data Fig. 2).

We also asked whether our three single unit stability metrics were predictive of the stability of a single unit’s evoked response to a panel of odorants (Extended Data Fig. 1B-D, bottom and 1F-H, bottom). To estimate the similarity of a single unit’s odor responses across days we computed for each single unit a vector whose elements consisted of the spontaneous baseline-subtracted, trial-averaged firing rate during the stimulus epoch for each odorant. We then measured the Pearson’s correlation between these vectors for each single unit across pairs of days.

To measure the dependence of waveform similarity and centroid displacement on the similarity in ACG shape (Extended Data Fig. 4), we computed the Pearson’s correlation of ACG similarity with either waveform similarity or centroid displacement for all pairs of single units that were stable across a given interval on all days within that interval.

### Peristimulus time histograms

To visualize the firing of individual single units we computed a peristimulus time histogram (PSTH) across all trials for a given odorant stimulus on a given day, smoothed using a Gaussian kernel (*σ* = 0.4 sec, Fig. 1G, Extended Data Fig. 5). To visualize the activity of populations of single units we *z*-scored the smoothed PSTH by subtracting for each single unit its mean spontaneous baseline firing rate across all trials on a given day during the baseline epoch (4 sec prior to stimulus onset, ‘baseline firing rate’) and dividing by the standard deviation of firing rates during the baseline epoch. Thus, the heat maps (Fig. 2A, Fig. 3A) indicate changes in firing rate in units of standard deviation of spontaneous activity. These two methods were used for visualization purposes only; unless stated otherwise, all statistical analyses were performed on unsmoothed, spontaneous baseline-subtracted firing rate estimates.

### Linear classification

To assess the linear separability of population odor responses we constructed single-trial population vectors using baseline-subtracted firing rates integrated over four time bins (0 - 2 sec, 2 - 4 sec, 4 - 6 sec, 6 - 8 sec after the onset of the 4 sec odorant stimulus; time 0 is defined as the moment a logic pulse triggers the olfactometer’s four-way valve to route the odor stream to the nose port). We quantified responses during these four epochs to account for the broad diversity in temporal response profiles across single units (e.g. Fig. 1G, Extended Data Fig. 5), ranging from transient ON or OFF responses to sustained responses that last several seconds. Thus, each population vector was composed of N × 4 elements where N is the number of single units that were stable across the pair of comparison days. Analysis results are qualitatively similar across a range of numbers and widths of quantification bins.

These population vectors, in combination with odorant stimulus labels, were used to train a linear kernel L2-regularized support vector machine. For within-day classification we employed leave-one-out-cross-validation (LOOCV), training on all but one of the 8 × 7 = 56 trials on a given day (8 odorant stimuli, 7 trials each) and testing classifier prediction on the trial that was left out. This procedure was repeated until all trials on a given day were tested in this way. For across-day classification we trained a model on all 56 trials one day, and tested that model’s predictions on all 56 trials on another day. Performance on shuffled data was tested by randomly permuting the odorant stimulus labels on the test dataset. We did not stitch single units across experimental subjects; all classifier analyses were performed on datasets obtained from individual animals. To permit pooling of classifier performance results across animals, we were limited by the lowest number of stable single units for any across-day comparison in any animal (41 single units). In all comparisons for which there were > 41 stable single units we employed 100 randomly selected 41 single unit subsets.

When measuring classification performance during the first 200 ms of the odor response (Extended Data Fig. 9), the narrow temporal window provided relatively noisy spike rate estimates with which to train the classifier. We therefore optimized the analysis parameters to promote classification performance as follows. (1) We employed the dataset for which we had the highest minimum single unit yield across all intervals (from amongst those animals that were presented eight odorant stimuli). (2) We performed classification on only subsets of four out of the eight odors. Thus, performance was promoted by requiring only 4-way rather than 8-way classification. Moreover, this scheme permitted 70 permutations over the eight odor classes. (3) Finally, we employed *z*-scored estimated firing rates, rather than baseline-subtracted firing rates, to denoise the training data.

### Population response similarity and drift rate calculation

To measure the similarity of population responses over time, we computed the Pearson’s correlation between pairs of trial-averaged population vectors for each odor across all pairs of comparison days. Population vectors were computed as described above (see Linear classification), then averaged across all trials for across-day comparisons, or averaged across either even or odd trials for within-day comparisons. We then computed the Pearson’s correlation between pairs of trial-averaged population vectors (Fig. 2D-F).

To compute the rate of drift across days we first converted correlations across days to angles as follows,

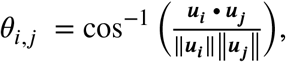

where *θ*_*i,j*_ is the angle between days *i* and *j* and ***u***_***i***_ is the trial-averaged population vector on day *i*. Variability within days was estimated as follows,

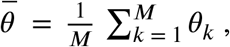

where *θ*_*k*_ is the mean angle across even and odd trials on each day for all days *k* out of *M* total days. We then estimated the rate of drift using two approaches. (1) We fit the trial-averaged population vector angles across all the intervals using a single exponential function : 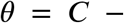 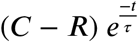, where *θ* is the variability (angle), *C* the asymptote, *R* the intercept at *t* = 0 (within-day variability, 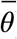), and *τ* the time constant of the exponential in days (Extended Data Fig. 8B). We then computed the drift rate over the 32-day interval by measuring the mean rate of change of the exponential fit. (2) We measured the drift rate across individual pairs of comparison days for individual odors as follows. For each odor, for each pair of days *i* and *j*, we compute the across-day angle *θ_i,j_* between trial-averaged population vectors. To correct for within-day variability, without which we would overestimate across-day drift^33^, we subtracted the within-day angle 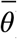. Drift rate (*r_i,j_*) across the interval between and *i*^th^ and *j*^th^ days was 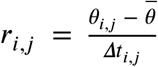 (Extended Data Fig. 8).

### Measurement of population response geometry

In order to assess whether the geometry of the population response in piriform drifted or remained the same across days, we first visualized population responses using principal component analysis (PCA). For this we computed the projection onto the first two principal components (PCs) of the trial averaged population vectors corresponding to 8 different odors. These PCs were computed separately on each individual day.

PCA was used for visualization purposes only; all statistical analyses were performed in the full dimensionality of the representations. For this, we first captured, for each day, the geometry of the population response by computing the odor-odor correlation matrix using trial-averaged population vectors. To measure how similar these geometries were across days, we computed the Frobenius norm between pairs of odor-odor correlation matrices. A Frobenius norm of 0 indicates identical matrices, whereas norms of increasing value indicate increasing difference between the matrices, and therefore increasing difference between the geometries encoded by those matrices. We computed an upper bound by computing this Frobenius norm across pairs of odor-odor correlation matrices in which the order of the trial-averaged population vectors was shuffled. These analyses were performed in individual animals (i.e. without stitching single units across animals) and only in those that were presented the panel of eight stimuli.

### Statistics

All statistical analyses were performed using custom software written in MATLAB (MathWorks, Natick, MA). Support vector machine analyses were implemented using LIBLINEAR^60^. Auto- and cross-correlograms were computed using spikes (github.com/cortex-lab/spikes). Except when otherwise noted we employed the nonparametric two-sided Wilcoxon rank-sum test to test the null hypothesis that the distribution of measurements in a pair of populations have equal medians.

To identify responsive odor-unit pairs (for Fig. 2B and Fig. 3B) we used a significance level of *α* = 0.001 (Wilcoxon rank-sum between the spike count during the 4 sec epoch before simulus onset for all trials on a given day and the spike count on the 7 trials during the odorant stimulus). Analysis results are qualitatively similar across different choices of *α*.

To quantify the fraction of single units that are stable across a 32-day interval, we computed binarized, signed tuning vectors for each single unit (0 if the single unit did not have a significant response to that odorant, 1 if the odorant evoked a significant increase in firing rate, −1 if the odorant evoked a significant decrease in firing rate, *α* = 0.001). We then defined a single unit as stable if its binarized tuning vectors were unchanged across the 32-day interval.

We measured the sparseness of the piriform population response across days using standard methods (Fig. 3B)^5,61–63^. The ‘population sparseness’ (*S_P_*) for each odorant was defined,

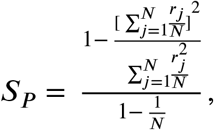

where *N* is the number of single units and *r_j_* is the response (spontaneous baseline-subtracted spike count during the 4 sec odor epoch) of the *j*^th^ single unit. We computed the ‘lifetime sparseness’ for each single unit using the same equation, but with *N* corresponding to the number of odorants tested and *r_j_* the response of a given single unit to the *j*^th^ odorant.

## References

1. Haberly, L. B. Single unit responses to odor in the prepyriform cortex of the rat. Brain Res. 12, 481–484 (1969).

2. Kadohisa, M. & Wilson, D. A. Separate encoding of identity and similarity of complex familiar odors in piriform cortex. Proc. Natl. Acad. Sci. 103, 15206–15211 (2006).

3. Rennaker, R. L., Chen, C.-F. F., Ruyle, A. M., Sloan, A. M. & Wilson, D. A. Spatial and temporal distribution of odorant-evoked activity in the piriform cortex. J. Neurosci. 27, 1534–1542 (2007).

4. Stettler, D. D. & Axel, R. Representations of odor in the piriform cortex. Neuron 63, 854–864 (2009).

5. Miura, K., Mainen, Z. F. & Uchida, N. Odor Representations in Olfactory Cortex: Distributed Rate Coding and Decorrelated Population Activity. Neuron 74, 1087–1098 (2012).

6. Bolding, K. A. & Franks, K. M. Recurrent cortical circuits implement concentration-invariant odor coding. Science 361, (2018).

7. Haberly, L. B. & Price, J. L. The axonal projection patterns of the mitral and tufted cells of the olfactory bulb in the rat. Brain Res. 129, 152–157 (1977).

8. Scott, J. W., McBride, R. L. & Schneider, S. P. The organization of projections from the olfactory bulb to the piriform cortex and olfactory tubercle in the rat. J. Comp. Neurol. 194, 519–534 (1980).

9. Nagayama, S. et al. Differential axonal projection of mitral and tufted cells in the mouse main olfactory system. Front. Neural Circuits 4, 120 (2010).

10. Sosulski, D. L., Bloom, M. L., Cutforth, T., Axel, R. & Datta, S. R. Distinct representations of olfactory information in different cortical centres. Nature 472, 213–216 (2011).

11. Ghosh, S. et al. Sensory maps in the olfactory cortex defined by long-range viral tracing of single neurons. Nature 472, 217–220 (2011).

12. Miyamichi, K. et al. Cortical representations of olfactory input by trans-synaptic tracing. Nature 472, 191–196 (2011).

13. Haberly, L. B. & Price, J. L. Association and commissural fiber systems of the olfactory cortex of the rat. I. Systems originating in the piriform cortex and adjacent areas. J. Comp. Neurol. 178, 711–740 (1978).

14. Johnson, D. M., Illig, K. R., Behan, M. & Haberly, L. B. New features of connectivity in piriform cortex visualized by intracellular injection of pyramidal cells suggest that “primary” olfactory cortex functions like “association” cortex in other sensory systems. J. Neurosci. 20, 6974–6982 (2000).

15. Franks, K. M. et al. Recurrent circuitry dynamically shapes the activation of piriform cortex. Neuron 72, 49–56 (2011).

16. Kaas, J. H. et al. Reorganization of retinotopic cortical maps in adult mammals after lesions of the retina. Science 248, 229–231 (1990).

17. Gilbert, C. D. & Wiesel, T. N. Receptive field dynamics in adult primary visual cortex. Nature 356, 150–152 (1992).

18. Keck, T. et al. Massive restructuring of neuronal circuits during functional reorganization of adult visual cortex. Nat. Neurosci. 11, 1162–1167 (2008).

19. Poort, J. et al. Learning enhances sensory and multiple non-sensory representations in primary visual cortex. Neuron 86, 1478–1490 (2015).

20. Rose, T., Jaepel, J., Hübener, M. & Bonhoeffer, T. Cell-specific restoration of stimulus preference after monocular deprivation in the visual cortex. Science 352, 1319–1322 (2016).

21. Clark, S. A., Allard, T., Jenkins, W. M. & Merzenich, M. M. Receptive fields in the body-surface map in adult cortex defined by temporally correlated inputs. Nature 332, 444–445 (1988).

22. Margolis, D. J. et al. Reorganization of cortical population activity imaged throughout long-term sensory deprivation. Nat. Neurosci. 15, 1539–1546 (2012).

23. Mayrhofer, J. M., Haiss, F., Helmchen, F. & Weber, B. Sparse, reliable, and long-term stable representation of periodic whisker deflections in the mouse barrel cortex. Neuroimage 115, 52–63 (2015).

24. Weinberger, N. M., Javid, R. & Lepan, B. Long-term retention of learning-induced receptive-field plasticity in the auditory cortex. Proc. Natl. Acad. Sci. 90, 2394–2398 (1993).

25. Recanzone, G. H., Schreiner, C. E. & Merzenich, M. M. Plasticity in the frequency representation of primary auditory cortex following discrimination training in adult owl monkeys. J. Neurosci. 13, 87–103 (1993).

26. Kato, H. K., Gillet, S. N. & Isaacson, J. S. Flexible sensory representations in auditory cortex driven by behavioral relevance. Neuron 88, 1027–1039 (2015).

27. Mombaerts, P. et al. Visualizing an olfactory sensory map. Cell 87, 675–686 (1996).

28. Kato, H. K., Chu, M. W., Isaacson, J. S. & Komiyama, T. Dynamic sensory representations in the olfactory bulb: modulation by wakefulness and experience. Neuron 76, 962–975 (2012).

29. Poo, C. & Isaacson, J. S. Odor representations in olfactory cortex:“sparse” coding, global inhibition, and oscillations. Neuron 62, 850–861 (2009).

30. Tolias, A. S. et al. Recording chronically from the same neurons in awake, behaving primates. J. Neurophysiol. 98, 3780–3790 (2007).

31. Dickey, A. S., Suminski, A., Amit, Y. & Hatsopoulos, N. G. Single-unit stability using chronically implanted multielectrode arrays. J. Neurophysiol. 102, 1331–1339 (2009).

32. Dhawale, A. K. et al. Automated long-term recording and analysis of neural activity in behaving animals. Elife 6, e27702 (2017).

33. Chestek, C. A. et al. Single-neuron stability during repeated reaching in macaque premotor cortex. J. Neurosci. 27, 10742–10750 (2007).

34. Pashkovski, S. L. et al. Structure and flexibility in cortical representations of odour space. Nature 583, 253–258 (2020).

35. Uchida, N. & Mainen, Z. F. Speed and accuracy of olfactory discrimination in the rat. Nat. Neurosci. 6, 1224–1229 (2003).

36. Wilson, C. D., Serrano, G. O., Koulakov, A. A. & Rinberg, D. A primacy code for odor identity. Nat. Commun. 8, 1–10 (2017).

37. Fink, A. J., Axel, R. & Schoonover, C. E. A virtual burrow assay for head–fixed mice measures habituation, discrimination, exploration and avoidance without training. Elife 8, e45658 (2019).

38. Fusi, S., Drew, P. J. & Abbott, L. F. Cascade models of synaptically stored memories. Neuron 45, 599–611 (2005).

39. Roxin, A. & Fusi, S. Efficient partitioning of memory systems and its importance for memory consolidation. PLoS Comput Biol 9, e1003146 (2013).

40. McClelland, J. L., McNaughton, B. L. & O’Reilly, R. C. Why there are complementary learning systems in the hippocampus and neocortex: insights from the successes and failures of connectionist models of learning and memory. Psychol. Rev. 102, 419 (1995).

41. Squire, L. R., Genzel, L., Wixted, J. T. & Morris, R. G. Memory consolidation. Cold Spring Harb. Perspect. Biol. 7, a021766 (2015).

42. Káli, S. & Dayan, P. Off-line replay maintains declarative memories in a model of hippocampal-neocortical interactions. Nat. Neurosci. 7, 286–294 (2004).

43. Rule, M.E., O’Leary, T. & Harvey, C.D. Causes and consequences of representational drift. Curr. Opin. Neurobiol. 58, 141–147 (2019).

44. Rule, M. E. et al. Stable task information from an unstable neural population. Elife 9, e51121 (2020).

45. Brette, R. Is coding a relevant metaphor for the brain? Behav. Brain Sci. 42, (2019).

46. Price, J. L. An autoradiographic study of complementary laminar patterns of termination of afferent fibers to the olfactory cortex. J. Comp. Neurol. 150, 87–108 (1973).

47. Kentros, C. G., Agnihotri, N. T., Streater, S., Hawkins, R. D. & Kandel, E. R. Increased attention to spatial context increases both place field stability and spatial memory. Neuron 42, 283–295 (2004).

48. Mankin, E. A. et al. Neuronal code for extended time in the hippocampus. Proc. Natl. Acad. Sci. 109, 19462–19467 (2012).

49. Rubin, A., Geva, N., Sheintuch, L. & Ziv, Y. Hippocampal ensemble dynamics timestamp events in long-term memory. Elife 4, e12247 (2015).

50. Lee, J. S., Briguglio, J., Romani, S. & Lee, A. K. The statistical structure of the hippocampal code for space as a function of time, context, and value. bioRxiv 615203 (2019).

51. Driscoll, L. N., Pettit, N. L., Minderer, M., Chettih, S. N. & Harvey, C. D. Dynamic reorganization of neuronal activity patterns in parietal cortex. Cell 170, 986–999 (2017).

52. Galván, V. V., Chen, J. & Weinberger, N. M. Long-term frequency tuning of local field potentials in the auditory cortex of the waking guinea pig. JARO J. Assoc. Res. Otolaryngol. 2, 199 (2001).

53. Mank, M. et al. A genetically encoded calcium indicator for chronic in vivo two-photon imaging. Nat. Methods 5, 805 (2008).

54. Jeon, B. B., Swain, A. D., Good, J. T., Chase, S. M. & Kuhlman, S. J. Feature selectivity is stable in primary visual cortex across a range of spatial frequencies. Sci. Rep. 8, 1–14 (2018).

55. White, E. L. Thalamocortical synaptic relations: a review with emphasis on the projections of specific thalamic nuclei to the primary sensory areas of the neocortex. Brain Res. Rev. 1, 275–311 (1979).

## Supplementary References

56. Cui, X., Wiler, J., Dzaman, M., Altschuler, R. A. & Martin, D. C. In vivo studies of polypyrrole/peptide coated neural probes. Biomaterials 24, 777–787 (2003).

57. Ludwig, K. A. et al. Poly (3, 4-ethylenedioxythiophene)(PEDOT) polymer coatings facilitate smaller neural recording electrodes. J. Neural Eng. 8, 014001 (2011).

58. PyDAQmx : a Python interface to the National Instruments DAQmx driver, Pierre CLADÉ, http://pythonhosted.org/PyDAQmx/.

59. Pachitariu, M., Steinmetz, N. A., Kadir, S. N., Carandini, M. & Harris, K. D. Fast and accurate spike sorting of high-channel count probes with KiloSort. in Advances in neural information processing systems 4448–4456 (2016).

60. Fan, R.-E., Chang, K.-W., Hsieh, C.-J., Wang, X.-R. & Lin, C.-J. LIBLINEAR: A library for large linear classification. J. Mach. Learn. Res. 9, 1871–1874 (2008).

61. Perez-Orive, J. et al. Oscillations and sparsening of odor representations in the mushroom body. Science 297, 359–365 (2002).

62. Rolls, E. T. & Tovee, M. J. Sparseness of the neuronal representation of stimuli in the primate temporal visual cortex. J. Neurophysiol. 73, 713–726 (1995).

63. Willmore, B. & Tolhurst, D. J. Characterizing the sparseness of neural codes. Netw. Comput. Neural Syst. 12, 255–270 (2001).

